# Anthrax Toxin as a Molecular Platform to Target Nociceptive Neurons and Modulate Pain

**DOI:** 10.1101/2020.03.28.004150

**Authors:** Nicole J. Yang, Jörg Isensee, Dylan Neel, Sai Man Liu, Han Xiong Bear Zhang, Andreea Belu, Shilpa Palan, Angela Kennedy-Curran, Pascal Röderer, Anja Nitzsche, Mike Lu, Bradley L. Pentelute, Oliver Brüstle, Vineeta Tripathi, Jin Mo Park, R. John Collier, Keith A. Foster, Bruce P. Bean, Stephen H. Leppla, Tim Hucho, Isaac M. Chiu

**Affiliations:** Department of Immunology, Harvard Medical School, Boston, MA 02115, USA; Translational Pain Research, Department of Anesthesiology and Intensive Care Medicine, University Hospital of Cologne, University of Cologne, 50931 Cologne, Germany; Ipsen Bioinnovation Ltd, Milton Park, Abingdon, OX14 4RY, UK; Department of Neurobiology, Harvard Medical School, Boston, MA 02115, USA; LIFE & BRAIN GmbH, Cellomics Unit, 53127 Bonn, Germany; Institute of Reconstructive Neurobiology, LIFE & BRAIN Center, Medical Faculty, University of Bonn, 53127 Bonn, Germany; Department of Chemistry, Massachusetts Institute of Technology, Cambridge, MA 02139, USA; Cutaneous Biology Research Center, Massachusetts General Hospital and Harvard Medical School, Charlestown, MA 02129, USA; Department of Microbiology, Harvard Medical School, Boston, MA 02115, USA; Laboratory of Parasitic Diseases, National Institute of Allergy and Infectious Diseases, National Institutes of Health, Bethesda, MD 20892, USA

**Author notes:** Correspondence to: Isaac Chiu Harvard Medical School Department of Immunology 77 Avenue Louis Pasteur Boston, MA 02115.

## Abstract

Bacterial toxins are able to act on neurons to modulate signaling and function. Here, we find that nociceptive sensory neurons that mediate pain are enriched in the receptor for anthrax toxins, ANTXR2. Anthrax Edema Toxin (ET) induced cAMP and PKA signaling in Nav1.8^+^ nociceptive neurons and modulated pain *in vivo*. Peripherally administered ET mediated mechanical allodynia in naïve mice and during *B. anthracis* infection. Intrathecally administered ET produced analgesic effects, potently blocking pain-like behaviors in multiple mouse models of inflammatory and chronic neuropathic pain. Nociceptor-specific ablation of ANTXR2 attenuated ET-induced signaling and analgesia. Modified anthrax toxin successfully delivered exogenous protein cargo into nociceptive neurons, illustrating utility of the anthrax toxin system as a molecular platform to target pain. ET further induced signaling in human iPSC-derived sensory neurons. Our findings highlight novel interactions between a bacterial toxin and nociceptors that may be utilized for developing new pain therapeutics.

**SUMMARY:** ANTXR2 expression on nociceptive neurons allows selective targeting and modulation of pain by native and engineered anthrax toxins.

## INTRODUCTION

Pain is an unpleasant sensation that is initiated by peripheral nociceptive neurons which detect intense thermal, mechanical and chemical stimuli *(1)*. Afferent nociceptive neurons have cell bodies that reside in the dorsal root ganglia (DRG) and transduce noxious signals from peripheral tissues to the spinal cord. Although acute pain is a critical protective mechanism that alerts the organism to danger, chronic pain is a pathological symptom that afflicts over 20% of the adult U.S. population *(2)*. However, existing analgesic therapies for chronic pain conditions are often insufficient or produce limiting side effects. For example, opioids, although widely prescribed, carry the risk of dependence, respiratory depression, constipation and nausea due to their off-target effects on the central nervous system (CNS) and enteric nervous system (ENS) *(3)*. Ongoing efforts to develop improved pain medications highlight the need for strategies to selectively target and silence nociceptive neurons and pain circuits.

Naturally occurring biological toxins are a rich source of evolutionary selected molecular agents that target neuronal function. In recent years, appreciation has grown for the range of bacterial products that affect the nervous system during homeostasis or pathogenesis *(4)*. We and others have found that specific microbial products can act on nociceptive neurons to modulate pain during pathogenic infection, and that nociceptive neurons can modulate host immune defenses. Here, we aimed to identify receptors for bacterial products expressed on nociceptive neurons and determine whether functional interactions between these microbial factors and neurons can regulate pain. By mining transcriptional data, we identified ANTXR2, the high affinity receptor for anthrax toxin, to be enriched in nociceptors. This observation suggested that anthrax toxin may target these neurons.

*Bacillus anthracis* is a gram-positive, sporulating bacterium that mainly affects grazing animals but can also cause cutaneous, gastrointestinal, respiratory or systemic infections in humans. Anthrax toxin is a major virulence factor of *B. anthracis*, composed of three proteins that come together to form two bipartite toxins: Lethal Toxin (LT), composed of Protective Antigen (PA) and Lethal Factor (LF), and Edema Toxin (ET), composed of PA and Edema Factor (EF). The intoxication mechanisms of anthrax toxin are well characterized, where PA first binds to cell surface anthrax toxin receptors, becoming proteolytically activated by furin-like proteases to oligomerize into a pore *(5)*. LF and EF bind to the PA pore through their N terminal domains LFN and EFN, and translocate into the cytoplasm where they are enzymatically active. LF is a zinc metalloprotease that cleaves MAP kinase kinases (MAPKKs) *(6)*, whereas EF is a calcium- and calmodulin-dependent adenylyl cyclase that converts ATP into cAMP *(7)*. The toxins disable the innate immune system at early stages of infection *(8)*, then directly damage tissue at later stages *(9)*.

There are two validated receptors for PA, which are structurally homologous: ANTXR1 (also known as TEM8) and ANTXR2 (also known as CMG2) *(10, 11)*. PA binds to ANTXR2 with higher affinity than ANTXR1 *(12, 13)*. ANTXR2-deficient mice but not ANTXR1-deficient mice are resistant to challenge by anthrax toxins and *B. anthracis* infection in survival and pathology *(14)*. These observations show that ANTXR2 is the major, physiologically relevant receptor for anthrax toxin *in vivo*.

Beyond their role in pathogenesis, anthrax toxins have been utilized as a delivery system for transporting functional cargo into the cytoplasm. Linking LFN with exogenous molecules has been shown to enable their delivery through the PA pore into target cells. Cargos that have been delivered include protein toxins *(15, 16)*, protein binders *(17)*, non-canonical polypeptides *(18)* and nucleic acids *(19)*. The breadth of these applications underscores the flexibility of the anthrax toxin system and its usefulness as a potential research and therapeutic tool.

Here, we describe a striking pattern of ANTXR2 expression in the nervous system, where it is mostly absent in CNS neurons but enriched in nociceptive DRG neurons. Intraplantar administration of ET produced hyperalgesia, whereas intrathecal administration of ET produced potent analgesia. We found that intrathecally administered ET targeted DRG neurons to induce cAMP and PKA signaling, and effectively blocked pain across multiple mouse models. Anthrax toxin also successfully delivered multiple types of non-native cargo into nociceptive neurons, demonstrating its potential as a broader platform to modulate nociception and pain. Altogether, we identify ANTXR2 as a marker enriched in nociceptive neurons and propose novel strategies for silencing pain based on anthrax toxin-mediated targeting.

## RESULTS

### ANTXR2 is expressed in Nav1.8^+^ nociceptive neurons and absent from most CNS neurons

Pain signaling is mediated by nociceptive sensory afferent neurons, whose cell bodies reside in the dorsal root ganglia (DRG) and trigeminal ganglia (TG). Previously, we found that DRG neurons can respond to bacteria-derived molecular mediators such as pore-forming toxins and formylated peptides, leading to pain during infection *(20–22)*.

To determine novel mechanisms that mediate neuronal responses to bacterial products and potentially modulate pain, we mined transcriptome data of DRG neuron populations *(23)* to identify cell surface receptors that could respond to bacterial molecules. We found that *Antxr2*, the high affinity receptor for anthrax toxin, was enriched by 5-fold in Nav1.8 lineage (*Nav1.8-cre ^Rosa26-Tdtomato^*^+^) sensory neurons compared to Parvalbumin lineage (*Pvalb-cre^Rosa26-Tdtomato^*^+^) proprioceptor neurons (**Fig. 1a**). Nav1.8^+^ sensory neurons have been shown to mediate mechanical, cold, and inflammatory pain *(24)*. *Antxr2* was expressed at similar levels in both non-peptidergic isolectin B4 (IB4)^+^Nav1.8^+^ neurons and IB4^-^Nav1.8^+^ neurons compared to Parv^+^ neurons (**Fig. S1a**). We further examined published single cell transcriptome data of the mouse nervous system *(25)*, and found that expression of *Antxr2* correlated with that of *Scn10a* (Nav1.8) but not *Pvalb* in DRG neuron subsets (**Fig. 1b**). The *Antxr2* expressing subset also encompassed nociceptor-like markers including *Trpv1*, *P2rx3* and *Calca*, but not markers of C-LTMRs such as *Th*.

**Fig. 1:**
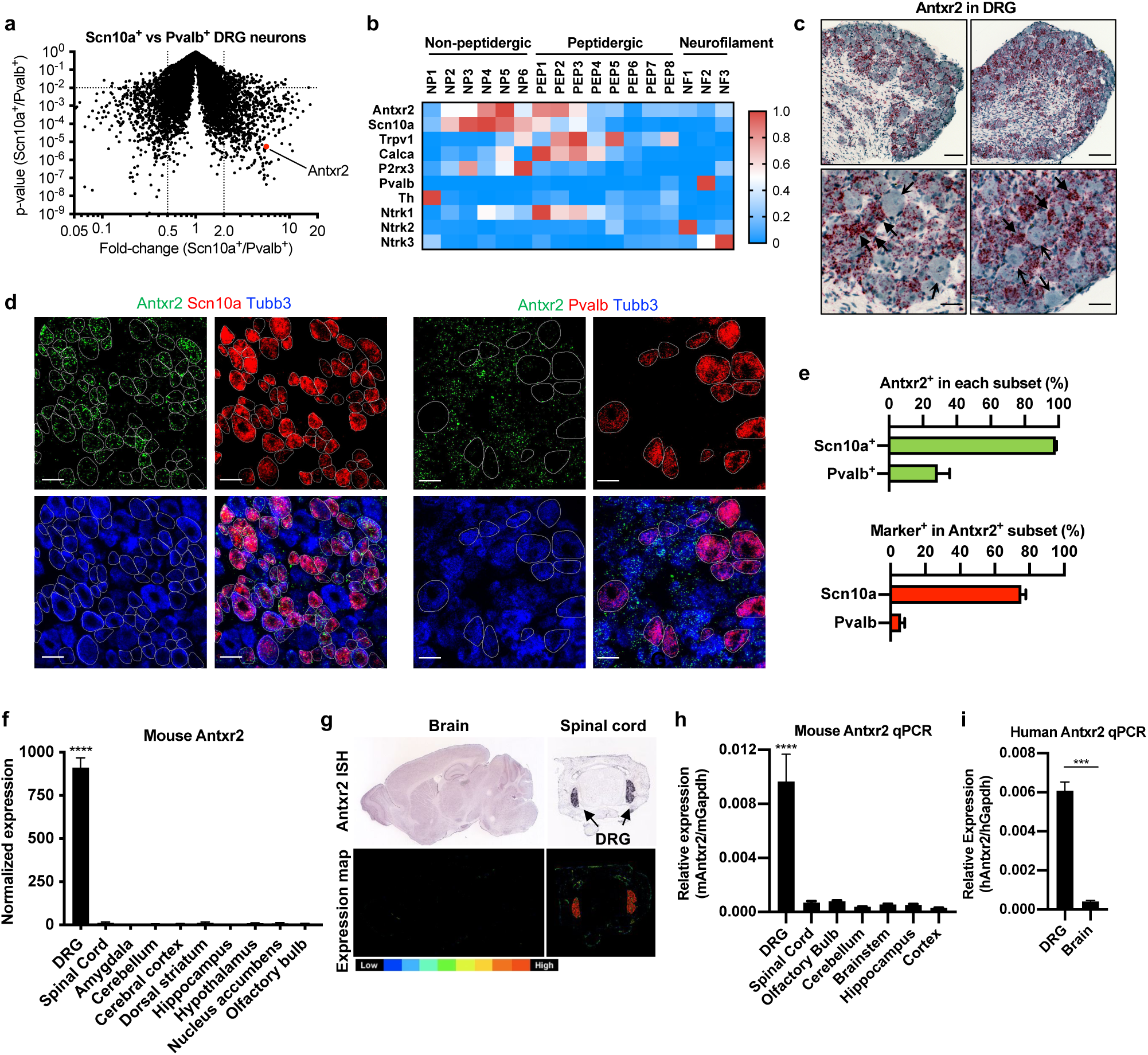
*Antxr2* is expressed in Nav1.8^+^ sensory neurons and absent from most CNS neurons. **(a)** Microarray analysis of preferentially expressed transcripts in sorted Nav1.8^+^ nociceptive neurons and Parv^+^ proprioceptive neurons *(23)*. **(b)** Expression of *Antxr2* and subgroup markers across sensory neuron subsets from published single cell RNA-seq data of the mouse nervous system *(25)*. **(c)** Representative images of *in situ* hybridization (ISH) analysis of *Antxr2* in the DRG. Solid and open arrowheads point to neurons with high or low levels of *Antxr2* transcripts, respectively. Scale bar, 100 μm (top row) or 35 μm (bottom row). **(d)** Representative images of multiplexed *in situ* hybridization (ISH) analysis of *Antxr2, Scn10a* and *Tubb3* (left) or *Antxr2, Pvalb* and *Tubb3* (right) in the DRG. Scale bar, 40 μm. **(e)** Quantification of multiplexed ISH images (n = 12-15 fields from 3 mice). Expression of *Antxr2*, *Scn10a* or *Pvalb* were scored in *Tubb3*^+^ neurons. **(f)** *Antxr2* expression in the DRG and brain regions from public microarray data *(97, 98)*. **(g)** *In situ* hybridization of *Antxr2* in adult brain (P56) and juvenile spinal cord (P4) from the Allen Mouse Brain Atlas and Mouse Spinal Cord Atlas *(99)*. Bottom row, color map of expression levels. **(h)** Quantitative PCR analysis of *Antxr2* expression in mouse DRG and brain (n = 3). **(i)** Quantitative PCR analysis of *Antxr2* expression from human DRG RNA (pooled from 4 individuals) and total brain RNA (pooled from 21 individuals). **Statistical analysis: (f, h)** One-way ANOVA with Dunnett’s post-test, DRG vs. all other tissue. **(h)** Unpaired t-test. \*\*\**p* < 0.001, \*\*\*\**p* < 0.0001.

This pattern of expression was also in line with another recently published single cell transcriptome dataset of DRG neurons across development *(26)*, where *Antxr2* expression in adult mice was enriched in CGRP^+^ nociceptors and non-peptidergic nociceptors, but not in proprioceptors or C-LTMRs (**Fig. S1b**). By contrast, *Antxr1* was largely absent or expressed at low levels in DRG neurons (**Fig. S1b**). RNAscope analysis validated the presence of *Antxr2* transcripts in the DRG, where highest level of expression was observed in small- and medium-diameter neurons (**Fig. 1c**). Large-diameter neurons, satellite glia and Schwann cells displayed low levels of expression. Quantification revealed that most *Scn10a*^+^ cells expressed *Antxr2*, whereas only a subset of *Pvalb*^+^ cells did (**Fig. 1d,e**), consistent with the transcriptional data. Conversely, most *Antxr2*-expressing neurons expressed *Scn10a*, whereas only a fraction expressed *Pvalb* (**Fig. 1e**). Collectively, our results demonstrated that *Antxr2* is expressed in somatosensory neurons and enriched in Nav1.8^+^ neurons.

While *Antxr2* is expressed in peripheral vascular and immune tissues, its presence across the nervous system has not been previously scrutinized. Mining tissue expression databases, we found that *Antxr2* is highly expressed in the mouse DRG but absent in CNS tissues including the spinal cord and multiple brain regions (**Fig. 1f**). We next analyzed *in situ* hybridization data from the Allen Brain Atlas. Strikingly, while *Antxr2* probe signal was strongly detected within the DRG, it was mostly absent throughout the brain and spinal cord (**Fig. 1g**). To confirm these results, we performed qPCR analysis for *Antxr2* expression in different mouse nervous tissues. We found that *Antxr2* expression is significantly higher in the DRG compared to various regions of the CNS, including the cortex, hippocampus, cerebellum, brainstem, olfactory bulb and spinal cord (**Fig. 1h**). We next examined *Antxr2* expression in human nervous tissue. Human *Antxr2* was expressed at higher levels in DRG RNA compared to brain RNA (**Fig. 1i**), consistent with our observations in mouse. Overall, we conclude that *Antxr2* is nearly absent from the brain and spinal cord but is enriched in Nav1.8^+^ sensory neurons in the DRG.

### Anthrax Lethal Toxin and Edema Toxin modulate signaling in nociceptive neurons

Given the enriched expression of *Antxr2* in DRG neurons, we next investigated whether Anthrax Lethal Toxin (LT) and Edema Toxin (ET) can functionally affect signaling. LT is composed of Protective Antigen (PA) and Lethal Factor (LF), where PA delivers LF into the cytoplasm of cells for cleavage of Map Kinase Kinases (MAPKKs). ET is composed of PA and Edema Factor (EF), where PA similarly delivers EF into the cytoplasm for generation of cAMP (**Fig. 2a**).

**Fig. 2:**
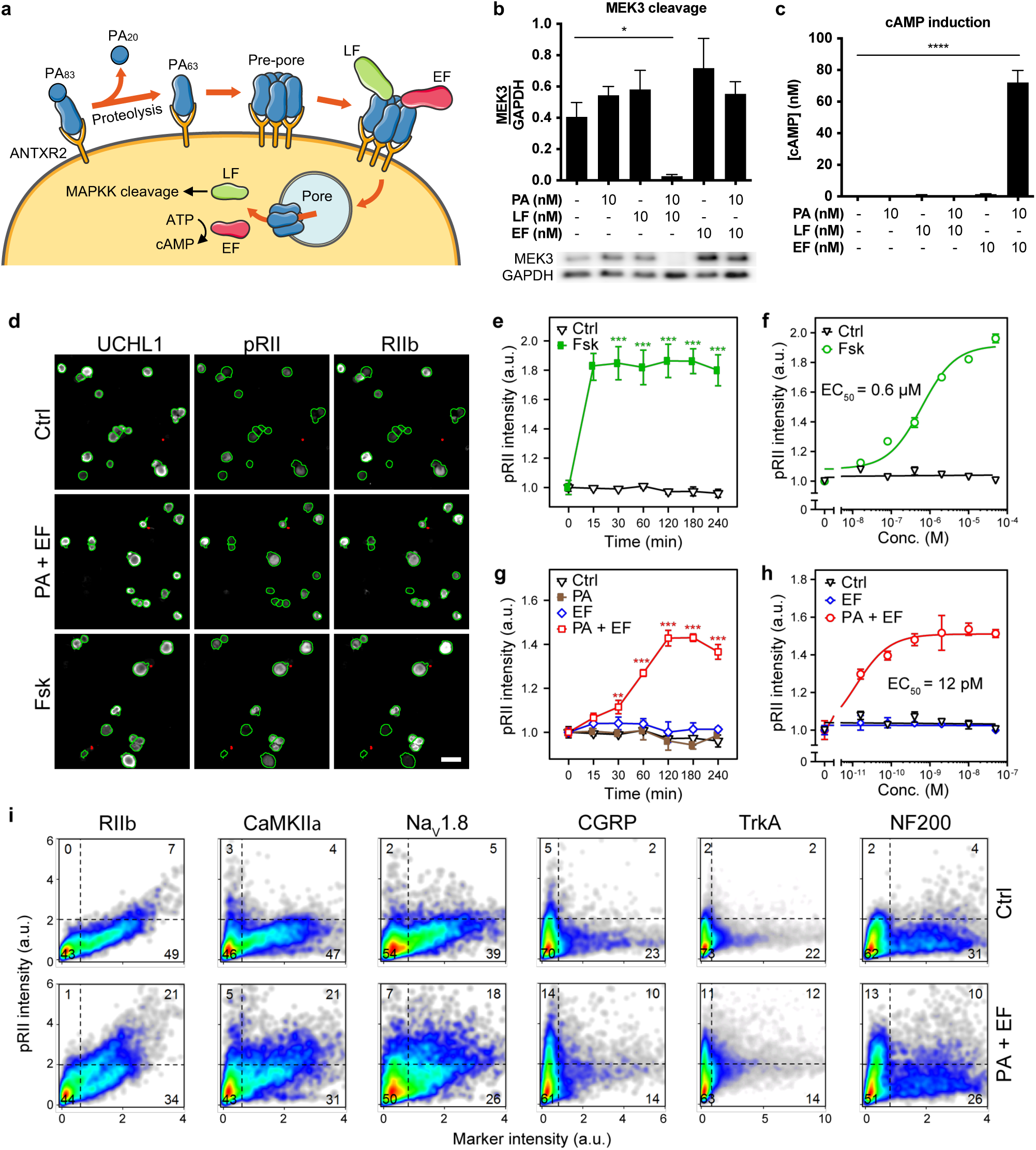
Anthrax Toxin modulates intracellular signaling in nociceptive neurons. **(a)** Schematic depicting the intoxication mechanism of anthrax lethal toxin (LT) and edema toxin (ET). **(b)** MEK3 expression levels measured by western blot in DRG cultures treated with PA, LF or EF for 24 h. Quantifications are from four independent experiments. **(c)** cAMP levels in DRG cultures treated with PA, LF or EF for 2 h. Results are from three independent experiments. **(d)** Representative HCS microscopy images of mouse DRG neurons stimulated with solvent control (Ctrl), EF (10 nM), or Fsk (10 µM) for 2 h. Cultures were immunolabeled with UCHL1 to identify the neurons, pRII to quantify PKA-II signaling activity, and RIIβ to identify nociceptors. Green or red encircled neurons indicate automatically selected or rejected objects, respectively (see Methods section). Scale bar, 50 µm. **(e)** Time-course of pRII intensity in DRG neurons stimulated with Fsk (10 µM). **(f)** Dose-response curve of pRII intensity in DRG neurons exposed to Fsk (0 - 50 µM, 2 h). **(g)** Time-course of pRII intensity in DRG neurons stimulated with Ctrl (0.1% BSA), PA (10 nM), EF (10 nM) or the combination of both factors. **(h)** Dose-response curve of pRII intensity in DRG neurons exposed to EF (0 - 50 nM, 2 h) in the presence of a constant dose of PA (10 nM). **(i)** Single cell data of DRG neurons stimulated with control solvent (0.1% BSA) or EF and PA (10 nM each) for 2 h. The DRG neurons were stained for UCHL1, pRII, and the indicated subgroup markers (>5000 neurons per marker). **Statistical analysis: (b, c)** One-way ANOVA with Dunnett’s post-test. **(e-h)** Error bars represent means ± SEM; *n* = 3 experiments; >2500 neurons/condition; two-way ANOVA with Bonferroni’s post-test. \**p* < 0.05, \*\**p* < 0.01, \*\*\**p* < 0.001, \*\*\*\**p* < 0.0001.

We first analyzed whether LT can modulate MAPK signaling in primary mouse DRG cultures. We treated DRG cultures with various combinations of anthrax toxin components (PA, LF, or EF) and probed for MEK3 cleavage and p38 phosphorylation. We observed significant cleavage of MEK3 (**Fig. 2b**) and reduced phosphorylation of p38 (**Fig. S2a**) specifically with LT (PA + LF) treatment. For example, LF alone had no effect on either measure, indicating that PA was required for delivery of LF into cells. EF with or without PA did not alter MEK3 levels.

We next tested whether ET can induce cAMP signaling in DRG cultures. We treated DRG cultures with combinations of PA, LF, or EF and subsequently measured cAMP levels in cell lysate following treatment. Significant increases in cAMP were observed only with ET (PA + EF) treatment, while PA alone, LF alone, or EF alone failed to have an effect (**Fig. 2c**). Furthermore, combined with a fixed concentration of PA (10 nM), EF potently induced cAMP with an EC_50_ of 48 pM (**Fig. S2b**). Our results show that anthrax toxins are able to act on DRG cultures to perturb intracellular signaling in a PA-dependent manner.

### Anthrax Edema Toxin modulates PKA signaling and excitability of nociceptive neurons

As our transcriptional analyses indicated that *Antxr2* is enriched in Nav1.8^+^ sensory neurons, we wished to functionally determine whether the anthrax toxins preferentially act on nociceptive neurons within DRG cultures. To this end, we utilized an automated High Content Screening (HCS) microscopy approach to quantify PKA activation in DRG neurons at the single cell level, in combination with neuronal marker stains *(27–30)*. PKA signaling is activated downstream of cAMP and is a key modulator of nociceptor signaling and pain *(31)*. As we previously demonstrated that ET elevates cAMP levels in DRG culture (**Fig. 2c** and **Fig. S2b**), we reasoned that the specific neurons that are targeted by PA to become intoxicated with EF could be identified by signs of PKA activation.

The type II isoforms of PKA (PKA-II) contain the regulatory subunit either RIIα or RIIβ (collectively referred to as RII), of which RIIβ is highly expressed in nociceptors *(27)*. The generation of cAMP by adenylyl cyclases such as EF triggers the dissociation of PKA catalytic subunits from regulatory subunits during kinase activation. In immunological staining of fixed neurons, the phosphorylated inhibitory site of RII (pRII) becomes accessible to antibodies after the catalytic subunits dissociate *(30, 32)*. We thus identified sensory neurons by ubiquitin C-terminal hydrolase L1 (UCHL1, formerly PGP9.5), nociceptors by RIIβ, and PKA-II activation by pRII (**Fig. 2d**).

First, we applied forskolin (Fsk) as a positive control to activate endogenous adenylyl cyclases in DRG neurons. Forskolin induced a long-lasting and dose-dependent increase in PKA-II activity in almost all DRG neurons (**Fig. 2e,f** and **Fig. S3a,b**). We next characterized the kinetics and dose dependence of ET treatment. Whereas stimulation with EF or PA alone had no effect, ET steadily increased PKA-II activity to saturate by 2 hours in a subgroup of DRG neurons (**Fig. 2g** and **Fig. S3a**). Stimulation with increasing doses of EF occurred with a picomolar EC_50_ value (measured at the 2 h time point; **Fig. 2h** and **Fig. S3b**), indicating highly effective uptake of ET. Stimulation with PA, LF or LT did not have any effect on PKA activation (**Fig. S3c, d**).

To characterize the subgroup of DRG neurons responding to ET, we stimulated DRG neurons with ET and stained the fixed cells for pRII and subgroup markers such as CaMKIIα and Nav1.8, markers for nociceptors; CGRP, a marker for peptidergic nociceptors; TrkA, a marker for nociceptive and thermoceptive sensory neurons; and NF200, a marker for large-diameter A-fiber neurons. We found that ET predominantly activated neurons expressing nociceptive markers such as RIIβ, CaMKIIα, Nav1.8, CGRP, and TrkA, rather than NF200-positive mechano- and proprioceptors (**Fig. 2i** and **Fig. S3e**). Overall, our analysis showed that ET dose-dependently targets nociceptive neuron subsets to activate PKA within hours of application.

PKA activation in nociceptors is known to enhance Nav1.8 *(33–36)* and HCN *(37, 38)* currents and reduce potassium currents *(39)*, resulting in hyperexcitability and hyperalgesia *(40)*. Our finding that ET activates PKA in nociceptive neurons raised the question of how ET affects their excitability. We hypothesized that ET may directly enhance neuronal excitability by inducing phosphorylation of voltage-gated channels. Alternatively, ET may deplete intracellular ATP, leading to activation of KATP channels, hyperpolarization, and reduced excitability. To test these two hypotheses, we examined the excitability of DRG neurons after ET treatment. DRG neurons were cultured overnight and then treated with ET (10 nM PA + 10 nM EF) or vehicle for at least 2 hours, after which neuronal excitability was tested. ET-treated neurons showed a substantial enhancement of excitability when tested by injection of small or moderate currents (**Fig. S4a-c**), consistent with the changes expected from activation of PKA. This enhancement of excitability occurred without detectable differences in resting membrane potential or input resistance of the neurons (**Fig. S4d-e**).

### Edema Toxin differentially modulates pain and cAMP signaling *in vivo*

Having determined that anthrax toxins act on nociceptive sensory neurons *in vitro*, we next examined whether the toxins affect pain behavior in mice. Both MAP kinase and cAMP pathways are implicated in nociceptor sensitization *(31)*, suggesting that anthrax toxin may potentially modulate pain. We first performed intraplantar injection of anthrax toxins into mice to determine if peripheral administration would alter pain-like behaviors. Mice were injected subcutaneously with combinations of PBS, PA alone, LT or ET. ET induced significant mechanical allodynia as assayed by von Frey filaments, which lasted for 8 hours post-injection (hpi) (**Fig. 3a**). ET additionally induced significant swelling of the footpad (**Fig. S5a**), as previously reported *(9)*. The degree of paw swelling did not strongly correlate with the degree of mechanical sensitization. Saline, PA alone, and LT had no effect on mechanical threshold sensitivities, indicating that ET has an acute effect on pain that the other toxins do not. To determine whether the activity of ET was confined to the periphery, we next measured cAMP levels in tissues following PBS or ET injection. A significant induction of cAMP at 2 hours post-injection of ET was observed in the footpad but not in the DRG or spinal cord (**Fig. 3b**). These results suggested that intraplantar ET induces local cAMP induction and pain hypersensitivity.

**Fig. 3:**
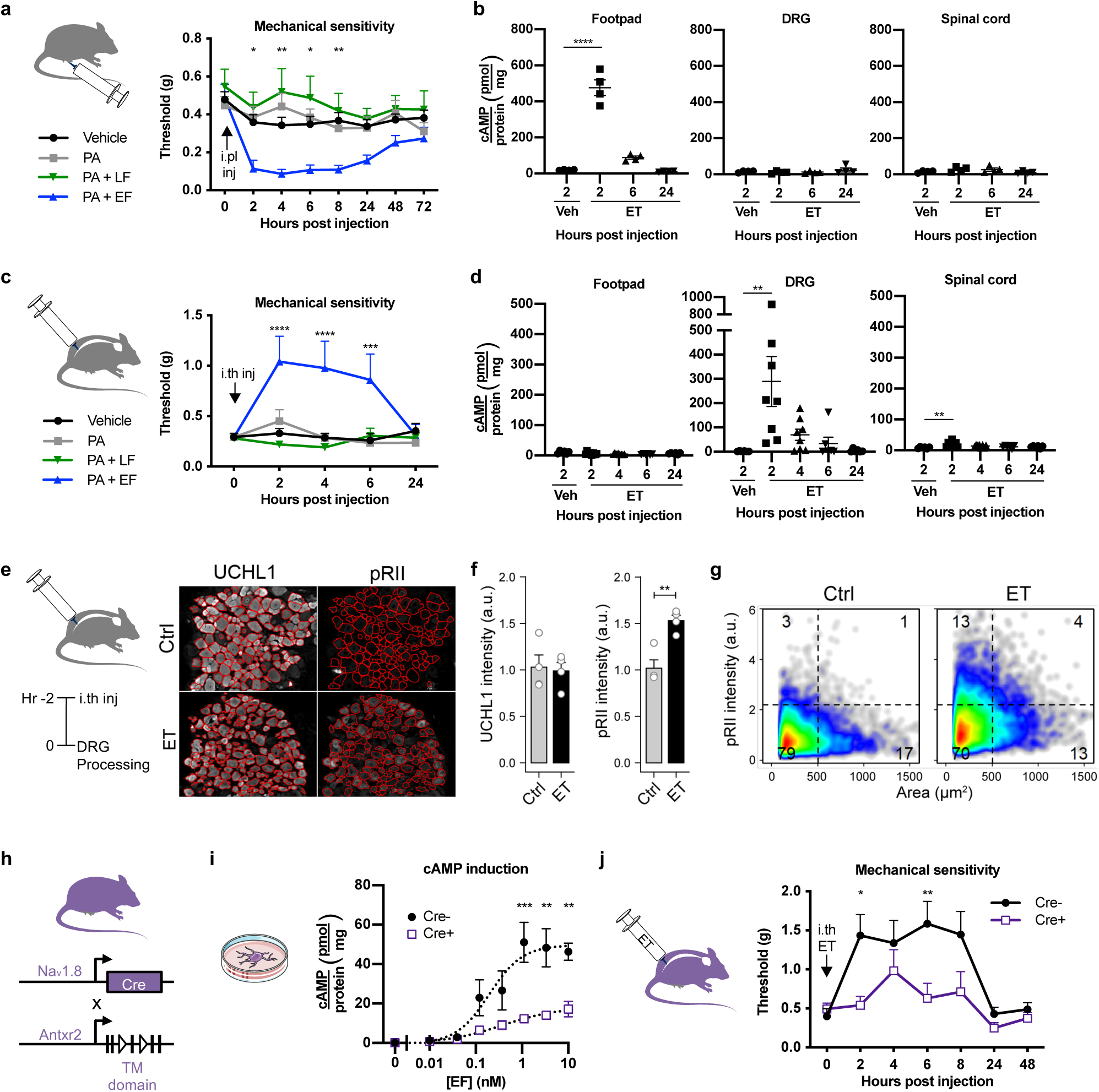
Edema Toxin modulates pain and cAMP signaling *in vivo*. **(a)** Mechanical sensitivity after intraplantar administration of vehicle (PBS), PA (2 μg), LT (2 μg PA + 2 μg LF) or ET (2 μg PA + 2 μg EF) (n = 9-17/group). **(b)** cAMP levels in the footpad, DRG or spinal cord after intraplantar administration of vehicle (PBS) or ET (2 μg PA + 2 μg EF) (n = 4/group). **(c)** Mechanical sensitivity after intrathecal administration of vehicle (PBS), PA (2 μg), LT (2 μg PA + 2 μg LF) or ET (2 μg PA + 2 μg EF) (n = 8/group). **(d)** cAMP levels in the footpad, DRG or spinal cord after intrathecal administration of vehicle (PBS) or ET (2 μg PA + 2 μg EF) (n = 6-8/group). **(e)** Representative images of frozen L3 - L6 DRG sections obtained from mice 2 h post intrathecal injection of vehicle (PBS) or ET (2 μg PA + 2 μg EF). The red lines indicate the mask used to quantify signal intensities in DRG neurons. **(f)** Mean UCHL1 and pRII intensities quantified in DRG sections of the respective mice (n = 4 animals per group, 15 - 20 images of 4 non-consecutive sections per animal, 1951 ± 279 neurons per animal). **(g)** Single cell data of the quantified DRG neurons. **(h)** Breeding strategy for generating conditional knock-out mice lacking functional ANTXR2 in Nav1.8 lineage nociceptors. **(i)** cAMP levels normalized to protein concentration in DRG neurons harvested from Nav1.8^cre/+^/Antxr2^fl/fl^ or Nav1.8^+/+^/Antxr2^fl/fl^ mice and treated with 10 nM PA and the indicated concentrations of EF for 2 h. **(j)** Mechanical sensitivity measured with von Frey filaments following intrathecal administration of ET (2 μg PA + 2 μg EF) to Nav1.8^cre/+^/Antxr2^fl/fl^ or Nav1.8^+/+^/Antxr2^fl/fl^ mice (n = 7/group). **Statistical analysis: (a, c)** Two-way ANOVA with Dunnett’s post-test, Vehicle vs. ET. **(b, d)** One-way ANOVA with Dunnett’s post-test. **(f)** Unpaired t-test. **(i, j)** Two-way ANOVA with Sidak’s post-test. \**p* < 0.05, \*\**p* < 0.01, \*\*\**p* < 0.001, \*\*\*\**p* < 0.0001.

Given the selectivity of ANTXR2 in the DRG compared to CNS tissues (**Fig. 1f-h**), we reasoned that intrathecal injections into the spinal subarachnoid space could better target anthrax toxins to DRG neurons while minimizing exposure to non-neuronal cells in the periphery. We thus next administered PA, LT or ET intrathecally to naïve mice. Surprisingly, we found that ET significantly raised mechanical sensitivity thresholds for several hours as assayed by von Frey filaments (**Fig. 3c**). Similarly, thermal pain thresholds assayed by the Hargreaves radiant heat test were elevated for 2 - 6 hpi (**Fig. S5b**). Unlike ET, intrathecal injections of LT, vehicle or PA alone did not affect mechanical or thermal thresholds, indicating that EF was required together with PA to induce these analgesic effects.

We next examined whether intrathecal ET induced signaling in specific regions of the nervous system or the periphery. We assayed for cAMP levels in the DRG, spinal cord, and footpad at different time points following vehicle or toxin administration. ET significantly increased cAMP levels in the DRG at 2 hpi, which gradually decreased by 4 - 6 hours and returned to baseline by 24 hours (**Fig. 3d**), an induction kinetics that mirrored the analgesic behavioral phenotype. In the spinal cord, cAMP induction was close to absent compared to the DRG with a minor increase at 2 hpi, and no changes were observed in the footpad. These results were consistent with selective expression of ANTXR2 in the DRG compared to the spinal cord. Intrathecal ET did not affect gross motor function and coordination as measured by the rotarod test (**Fig. S5c**). Although toxin administration slightly decreased retention times on the rotarod at 2 and 4 hpi, differences in magnitude were minor and not statistically significant.

Overall, our findings indicate that intraplantar administration of ET induces mechanical allodynia and local cAMP induction in the footpad, while intrathecal administration of ET induces analgesia and predominantly targets the DRG with minimal effects on the spinal cord.

### Intrathecal administration of ET activates PKA in neurons *in vivo*

Given that intrathecal but not intraplantar injection of ET elevated cAMP levels in the DRG, we next examined the effects of intrathecal ET injection on DRG neurons *in vivo*. We administered ET intrathecally to mice and used a semi-automated imaging and analysis approach to measure PKA-II activity in individual neurons within frozen sections of lumbar DRGs (L3 - L6) (**Fig. 3e**). The sections were stained for UCHL1 to identify neurons and pRII to measure PKA-II activation. Quantification of ≈2000 neurons per animal revealed substantial induction of PKA-II activity in DRG neurons with a similar magnitude as observed *in vitro* (**Fig. 3f, g**). To determine whether ET induces PKA-II activity in the central nociceptor terminals in the dorsal horn, we also analyzed corresponding lumbar spinal cord sections stained for UCHL1, RIIβ and pRII. As expected, RIIβ predominantly labelled the outer layers of the dorsal horn indicating the presence of nociceptor terminals (**Fig. S6a**). However, the pRII signal intensity in this region did not significantly differ with control or ET treatment (**Fig. S6b**), indicating that activation of PKA-II in central nociceptor terminals, if any, was not detectable above background. Altogether, our results demonstrated that intrathecally administered ET targets DRG neurons *in vivo* to activate cAMP-dependent signaling.

### Nociceptor neuron ablation of ANTXR2 attenuates Edema Toxin-induced signaling and analgesia

We next aimed to confirm whether the modulatory effects of ET on pain are mediated through targeting ANTXR2 on nociceptive neurons. To this end, we generated and characterized mice specifically lacking ANTXR2 function in nociceptors by breeding Nav1.8-cre knock-in mice *(24)* with a conditionally targeted allele of *Antxr2* in the transmembrane region (Antxr2^fl/fl^) *(9, 14)* (**Fig. 3h**). For subsequent experiments, these Nav1.8^cre/+^/Antxr2^fl/fl^ mice were compared to control Nav1.8^+/+^/Antxr2^fl/fl^ littermates. Reduced expression of the full-length *Antxr2* transcript was confirmed in the DRG (**Fig. S7a**).

Given that ANTXR2 is enriched in nociceptive neurons, we first determined whether deleting the receptor would affect pain behavior in the absence of toxin administration. The endogenous function of ANTXR2 is not fully understood, with reported roles in binding to extracellular matrix proteins and mediating their internalization and degradation *(41)*. Compared between littermates, the absence of ANTXR2 did not affect most pain modalities at baseline, including mechanical pain thresholds assayed with von Frey filaments; thermal pain thresholds assayed with the Hargreaves apparatus and hot plate; and cold plate responses (**Fig. S7b**). We observed a slight impairment in noxious mechanoception assayed with the Randall-Selitto test, which was statistically significant but minor in magnitude. Next, examining inflammatory and neuropathic pain behavior, we observed a slight decrease in the second phase of formalin-induced pain (**Fig. S7c**) and no defects in carrageenan-induced hyperalgesia (**Fig. S7d**). Overall our results indicated that ANTXR2 does not play a significant role in nociceptor function.

We next examined whether neuronal expression of ANTXR2 was relevant to ET-induced signaling and pain modulation. cAMP induction following ET treatment was significantly reduced in cultured DRG neurons harvested from Nav1.8^cre/+^/Antxr2^fl/fl^ mice (**Fig. 3i**). Next, we administered ET by intraplantar or intrathecal routes to Nav1.8^cre/+^/Antxr2^fl/fl^ mice and assayed for pain-like behavior. The hyperalgesia induced by intraplantar ET was unaffected in Nav1.8^cre/+^/Antxr2^fl/fl^ mice compared to littermate controls (**Fig. S7e**), suggesting that ET sensitizes nociceptors through an indirect mechanism in the periphery. By contrast, the analgesia induced by intrathecal ET was significantly attenuated in Nav1.8^cre/+^/Antxr2^fl/fl^ mice (**Fig. 3j**). Our results indicated that direct targeting of Nav1.8-expressing nociceptors via ANTXR2 contributes to the analgesia produced by intrathecal ET but not to the allodynia caused by intraplantar ET, which may occur through an indirect mechanism.

### Intrathecally administered Edema Toxin silences multiple modalities of neuropathic and inflammatory pain

Given that intrathecal injection of ET induced potent analgesia in naïve mice, we wondered whether it could be used as a broader molecular strategy to target nociceptive neurons and silence pain. We thus examined whether intrathecal administration of ET can silence different modalities of acute or chronic pain.

We first analyzed whether ET has an effect on the Spared Nerve Injury (SNI) model of neuropathic pain *(42)*. Mice underwent SNI, and 7 days post-surgery were intrathecally injected with saline or anthrax toxin components (PA, EF, ET or LT) when significant mechanical allodynia was established. ET effectively blocked SNI-induced mechanical allodynia for several hours, whereas PA alone, EF alone, or LT had no effect (**Fig. 4a**). Interestingly, ET raised the mechanical pain threshold in both the ipsilateral paw and contralateral paw of mice significantly above baseline (**Fig. 4b**). Correspondingly, cAMP levels in both the ipsilateral and contralateral DRGs (L3 – L5) were significantly elevated above vehicle at the same time points (**Fig. 4c**).

**Fig. 4:**
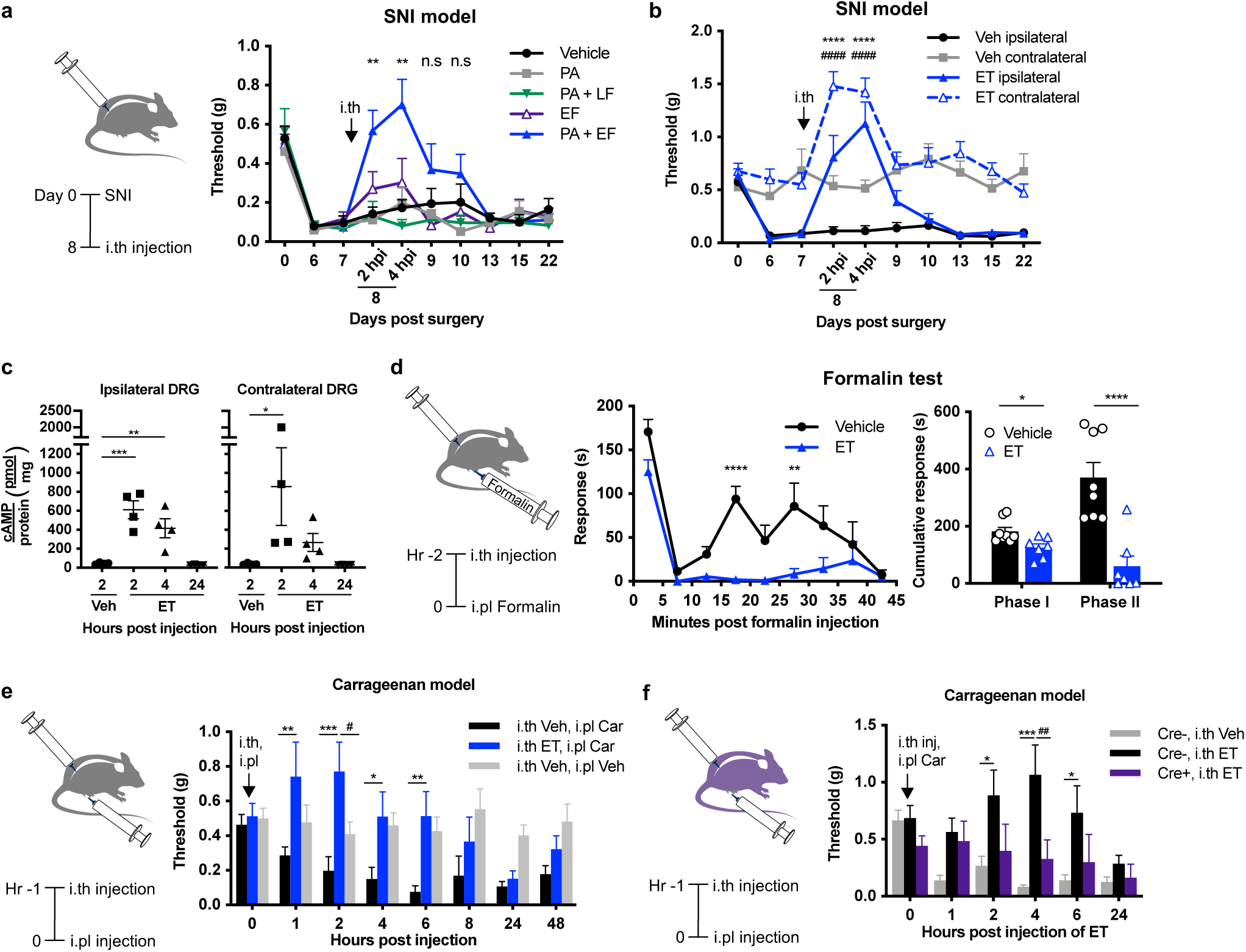
Intrathecally administered Edema Toxin silences multiple modalities of neuropathic and inflammatory pain. **(a)** Mechanical sensitivity in SNI mice following intrathecal administration of vehicle (PBS), PA (2 μg), LT (2 μg PA + 2 μg LF), EF (2 μg) or ET (2 μg PA + 2 μg EF) (n = 7-15/group). **(b)** Mechanical sensitivity in the ipsilateral or contralateral hind paw of SNI mice following intrathecal administration of vehicle (PBS) or ET (2 μg PA + 2 μg EF) (n = 9/group). **(c)** cAMP levels in the ipsilateral or contralateral DRGs (L3 – L5) of SNI mice following intrathecal administration of vehicle (PBS) or ET (2 μg PA + 2 μg EF) (n = 4/group). **(d)** Mice were given intrathecal injection of vehicle (PBS) or ET (2 μg PA + 2 μg EF) 2 h prior to intraplantar injection of 5% formalin. **(Left)** Acute pain behavior measured in 5 min intervals. **(Right)** Cumulative responses during Phase I (0 – 5 min) or Phase II (15 – 35 min) (n = 7-8/group). **(e)** Mechanical sensitivity measured with von Frey filaments. Mice were given intrathecal injection of vehicle (PBS) or ET (2 μg PA + 2 μg EF) 1 h prior to intraplantar injection of vehicle (0.9% saline) or 2% carrageenan (n = 8- 9/group). **(f)** Mechanical sensitivity measured with von Frey filaments. Nav1.8^cre/+^/Antxr2^fl/fl^ mice or Nav1.8^+/+^/Antxr2^fl/fl^ littermates were given intrathecal injection of vehicle (PBS) or ET (2 μg PA + 2 μg EF) 1 h prior to intraplantar injection of 2% carrageenan (n = 7-8/group). **Statistical analysis: (a)** Two-way ANOVA with Dunnett’s post-test. Vehicle vs. ET, \*\**p* < 0.01, n.s, not significant. **(b)** Two-way ANOVA with Tukey’s post-test. Vehicle ipsilateral vs. ET ipsilateral, \*\*\*\**p* < 0.0001. Vehicle contralateral vs. ET contralateral, ####*p* < 0.0001. **(c)** Two-way ANOVA with Dunnett’s post-test. **(d, left)** Two-way ANOVA with Sidak’s post-test. **(d, right)** Unpaired t-test. **(e)** Two-way ANOVA with Tukey’s post-test. i.th Vehicle, i.pl Carrageenan vs. i.th ET, i.pl Carrageenan, \**p* < 0.05, \*\**p* < 0.01, \*\*\**p* < 0.001. i.th Vehicle, i.pl Vehicle vs. i.th ET, i.pl Carrageenan, #*p* < 0.05. **(f)** Two-way ANOVA with Tukey’s post-test. Cre-, i.th Vehicle vs. Cre-, i.th ET, \**p* < 0.05, \*\*\**p* < 0.001. Cre-, i.th ET vs Cre+, i.th ET, ##*p* < 0.01.

We next analyzed the effects of ET in the formalin model of nocifensive pain. Administered intrathecally 2 hours prior to formalin, ET significantly decreased both the first and second phase of formalin-induced pain with notably strong effects in the second phase (**Fig. 4d**).

Finally, we tested ET intrathecal injection in the carrageenan model of inflammatory pain. ET but not vehicle injection produced significant attenuation of mechanical allodynia lasting for several hours (**Fig. 4e**). Notably at the 2 hours post-injection timepoint, mechanical pain thresholds were increased beyond that of controls which did not receive carrageenan, emphasizing the potent effects of ET. Furthermore, the analgesic effects of ET were attenuated in the carrageenan model in Nav1.8^cre/+^/Antxr2^fl/fl^ mice compared to littermate controls (**Fig. 4f**), indicating that ANTXR2-mediated targeting of Nav1.8 nociceptive neurons contributed to ET activity. Altogether, ET induced potent analgesia lasting for several hours in distinct models of inflammatory and neuropathic pain, suggesting it may act as a broadly applicable analgesic.

### Anthrax Edema Factor contributes to mechanical allodynia during subcutaneous *B. anthracis* infection

We next wished to determine whether the anthrax toxins contribute to pain in a physiologically relevant setting. The differential effects of ET on pain based on injection site (intraplantar vs. intrathecal) also led us to question whether infection by *B. anthracis* leads to pain or analgesia. To this end, we established a subcutaneous infection model in the footpad with vegetative *B. anthracis*, taking advantage of active toxin secretion by vegetative bacteria. We used the attenuated Sterne strain, which lacks the virulent capsule but expresses all components of anthrax toxin and is widely used for laboratory research *(43)*. Bacteria caused dose-dependent mortality over the course of 10 days (**Fig. S8a**), accompanied by significant swelling in the footpad and a dose-dependent increase in body weight (**Fig. S8b**). Bacterial infection also induced significant mechanical allodynia over the first 48 hours of infection, indicating development of pain hypersensitivity prior to the lethal time points of infection (**Fig. S8c**). To determine the role of Edema Factor in pain and infection, we compared mice infected with WT *B. anthracis* Sterne or with an isogenic mutant strain lacking edema factor (ΔEF). Deficiency in EF led to significantly increased survival, decreased local edema and body weight increases following infection (**Fig. S8d, e**). Mechanical allodynia was also significantly reduced in ΔEF infected mice compared to WT infected mice, indicating that EF contributes to the pain-like behaviors (**Fig. S8f**). These data show that EF contributes to the mortality, mechanical allodynia, edema and weight gain during infection. These findings may relate to the mechanical allodynia we observed with ET injection into the footpad (**Fig. 3a**).

### Anthrax toxin delivers exogenous molecular cargoes into neurons for functional modulation

The broad and potent analgesia induced by intrathecally administered ET suggested that the toxin could be utilized as an effective strategy to silence pain. Furthermore, ET efficiently targeted DRG neurons *in vivo* (**Fig. 3e-g**), supporting the use of PA as a selective binding moiety for ANTXR2-expressing nociceptors. We thus wondered whether the anthrax toxin system could be further expanded to deliver non-native cargo into nociceptive neurons and confer greater versatility in modulating function. The PA + LFN system based on anthrax lethal toxin has been widely demonstrated to deliver exogenous cargo into cancer cell lines and immune cells *(44, 45)*. Previously we had determined that LF can be delivered through PA to inhibit MAP kinase signaling in DRG cultures (**Fig. 2b**). We therefore reasoned that the existing PA + LFN system could be adapted to deliver molecular cargo into DRG neurons (**Fig. 5a**).

**Fig. 5:**
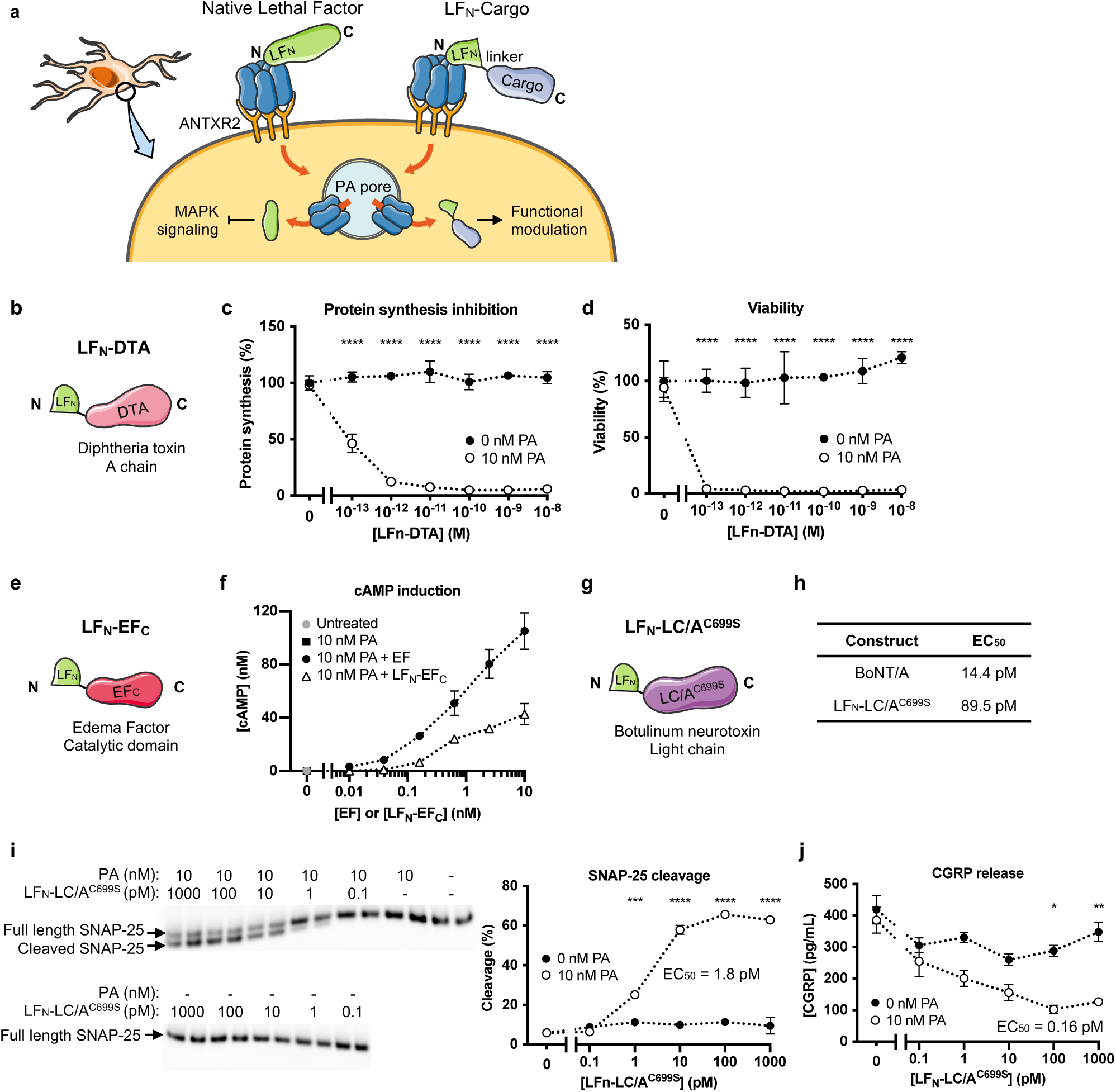
Anthrax toxin as a platform to deliver exogenous cargo into nociceptive neurons for functional modulation of activity. **(a)** Delivery of exogenous cargo into neurons by the PA + LFN system. **(b)** Design of LFN-DTA linking the N terminal domain of LF (LFN) to the A chain of diphtheria toxin (DTA). **(c)** Protein synthesis levels of DRG cultures following 6 h treatment with the indicated concentrations of LFN-DTA ± PA (10 nM). **(d)** Viability of DRG cultures following 7 day treatment with the indicated concentrations of LFN-DTA ± PA (10 nM). **(e)** Design of LFN-EFC linking the N terminal domain of LF (LFN) to the catalytic domain of EF (EFC). **(f)** cAMP levels in DRG cultures treated with 10 nM PA + EF or 10 nM PA + LFN-EFC for 2 h. **(g)** Design of LFN-LC/A^C699S^ linking the N terminal domain of LF (LFN) to a mutated light chain (LC) of type A botulinum neurotoxin (BoNT/A). **(h)** Enzymatic activity of BoNT/A or LFN-LC/A^C699S^ measured with a cell-free *in vitro* assay. **(i-j)** DRG cultures were treated with the indicated concentrations of LFN-LC/A^C699S^ ± PA (10 nM) for 24 h and stimulated with 80 mM KCl for 10 min. **(i)** SNAP-25 cleavage in cell lysate measured by western blot. Percent cleavage was calculated using band intensities with the following formula: cleaved/(intact + cleaved). **(j)** CGRP release in the supernatant. **Statistical analysis: (c, d, i, j)** Two-way ANOVA with Sidak’s post-test. \**p* < 0.05, \*\**p* < 0.01, \*\*\**p* < 0.001, \*\*\*\**p* < 0.0001.

To test this idea, we first utilized PA in combination with LFN-DTA, a widely used reagent for assaying PA-mediated internalization *(16)* (**Fig. 5b**). The A chain of Diphtheria Toxin (DTA) inactivates eukaryotic elongation factor 2 (eEF-2) to inhibit protein translation in mammalian cells, eventually causing cell death. We found that DRG cultures treated with PA and LFN-DTA potently blocked translation with sub-picomolar EC_50_ after 6 hours of treatment (**Fig. 5c**). Continued treatment led to significantly reduced viability by day 7 (**Fig. 5d**). LFN-DTA alone had no effect on both measures, demonstrating that internalization of LFN-DTA was specifically mediated by PA.

Next, we generated a chimera of LF and EF where LFN is linked to the C terminal catalytic domain of EF (EFC), referred to as LFN-EFC (**Fig. 5e**). We reasoned that successful delivery of EFC into nociceptors via the PA + LFN system could be confirmed by induction of cAMP. The N terminus of EF is homologous to LFN in sequence and structure *(46)*, and serves the same functional role of mediating binding to the PA pore and initiating translocation *(47)*. Cultured DRG neurons were treated with PA + LFN-EFC or PA + EF for comparison. For this experiment, we used an EF clone that was produced with the same *B. anthracis*-based expression system as LFN-EFC. In the presence of 10 nM PA, LFN-EFC robustly induced cAMP with an EC_50_ of 660 pM (**Fig. 5f**). Its potency and efficacy was slightly lower than EF, which may be due to the N terminal substitution impacting association with calmodulin *(48)*. There was no effect in the absence of PA, indicating that LFN-EFC enters the neurons specifically through PA pores.

Lastly, we designed and synthesized a novel LFN-based construct that can inhibit neurotransmitter release from nociceptive neurons. Botulinum neurotoxin (BoNT) cleaves components of the SNARE complex, preventing synaptic vesicles containing neurotransmitters from fusing with the plasma membrane. Structurally, BoNT is composed of two peptide chains linked with a disulfide bond, where the heavy chain (HC) mediates internalization of the enzymatic light chain (LC). We fused the light chain of BoNT serotype A1 (LC/A1), which targets SNAP-25, to the C terminus of LFN. In addition, a free cysteine towards the C terminus of LC/A1 was mutated to a serine to prevent the formation of intra- or inter-molecular disulfide bonds, leading to the final construct LFN-LC/A1^C699S^ (**Fig. 5g**).

LFN-LC/A1^C699S^ retained enzymatic activity in solution, with an EC_50_ of 89.5 pM measured in a cell-free assay (**Fig. 5h**). This EC_50_ was approximately six-fold higher than that of native BoNT/A (14.4 pM), but still potently active in the picomolar range. We next asked whether PA could deliver LFN-LC/A1^C699S^ into DRG neurons in culture to cleave SNAP-25 and inhibit the release of key neuropeptides. DRG neurons were cultured for one week to allow for expression of neuropeptides, then treated with varying concentrations of LFN-LC/A1^C699S^ in the presence or absence of PA. Combined with a fixed concentration of PA (10 nM), LFN-LC/A1^C699S^ produced dose-dependent cleavage of SNAP-25 with a picomolar EC_50_ (**Fig. 5i**). Release of the neuropeptide CGRP also showed a dose-dependent decrease with a sub-picomolar EC_50_ (**Fig. 5j**). Treatment with LFN-LC/A1^C699S^ alone or PA alone did not affect either SNAP-25 cleavage or CGRP release, suggesting that LFN-LC/A1^C699S^ is required and dependent on PA for cell entry.

Finally, we wished to test whether potential analgesic cargo can be delivered into sensory neurons in mice using the PA + LFN system. We first administered PA + LFN-LC/A1^C699S^ intrathecally to SNI mice and measured changes in mechanical pain thresholds, but did not observe any effects on pain. We hypothesized that stronger effects may be produced by driving greater accumulation of cargo through increased injections. As such, we reasoned that dosing the construct multiple times separated in time may allow ANTXR2 to be replenished on the cell surface to mediate multiple waves of internalization. Indeed, repeated intrathecal administration of PA + LFN-LC/A1^C699S^ performed on three consecutive days produced significant blockade of neuropathic pain in the SNI model (**Fig. S9**). Administration of PA or LFN-LC/A1^C699S^ alone had no effect, indicating that delivery of LFN-LC/A1^C699S^ was PA-mediated. Altogether, our results demonstrated that the PA + LFN system is capable of delivering molecularly and functionally distinct cargo into nociceptive neurons and targeting pain signaling *in vivo*, supporting further development as a molecular platform to target nociception and pain.

### Edema Toxin targets human iPSC-derived sensory neurons

As a potential approach to treat pain, we wished to test the relevance of anthrax toxin-based targeting strategies in human sensory neurons. We previously found that *Antxr2* is expressed in human DRGs (**Fig. 1i**), suggesting that anthrax toxin may functionally target human nociceptors. We thus applied our high content screening (HCS) microscopy approach to investigate whether ET activates cAMP signaling in human iPSC-derived sensory neurons. We differentiated human iPSCs into sensory neurons of a nociceptor phenotype following a previously established small molecule-based differentiation protocol *(49–52)*. After three weeks of maturation, iPSC-derived sensory neurons were stimulated with PA, EF or PA + EF, and fixed and stained for pRII (**Fig. 6a, b**). Stimulation with EF or PA alone did not activate PKA-II activity, whereas stimulation with PA + EF resulted in increased pRII activation (**Fig. 6c, d**). We next determined the potency of response by titrating the concentration of EF in the presence or absence of PA. At 2 hours, PA + EF produced a dose-dependent activation of PKA-II in human iPSC-derived sensory neurons with a picomolar EC_50_ value similar to that measured in the mouse *in vitro* system (**Fig. 2h**). These results support the susceptibility of human nociceptive neurons to anthrax toxin and underscore the potential of anthrax-toxin based strategies for silencing pain.

**Fig. 6:**
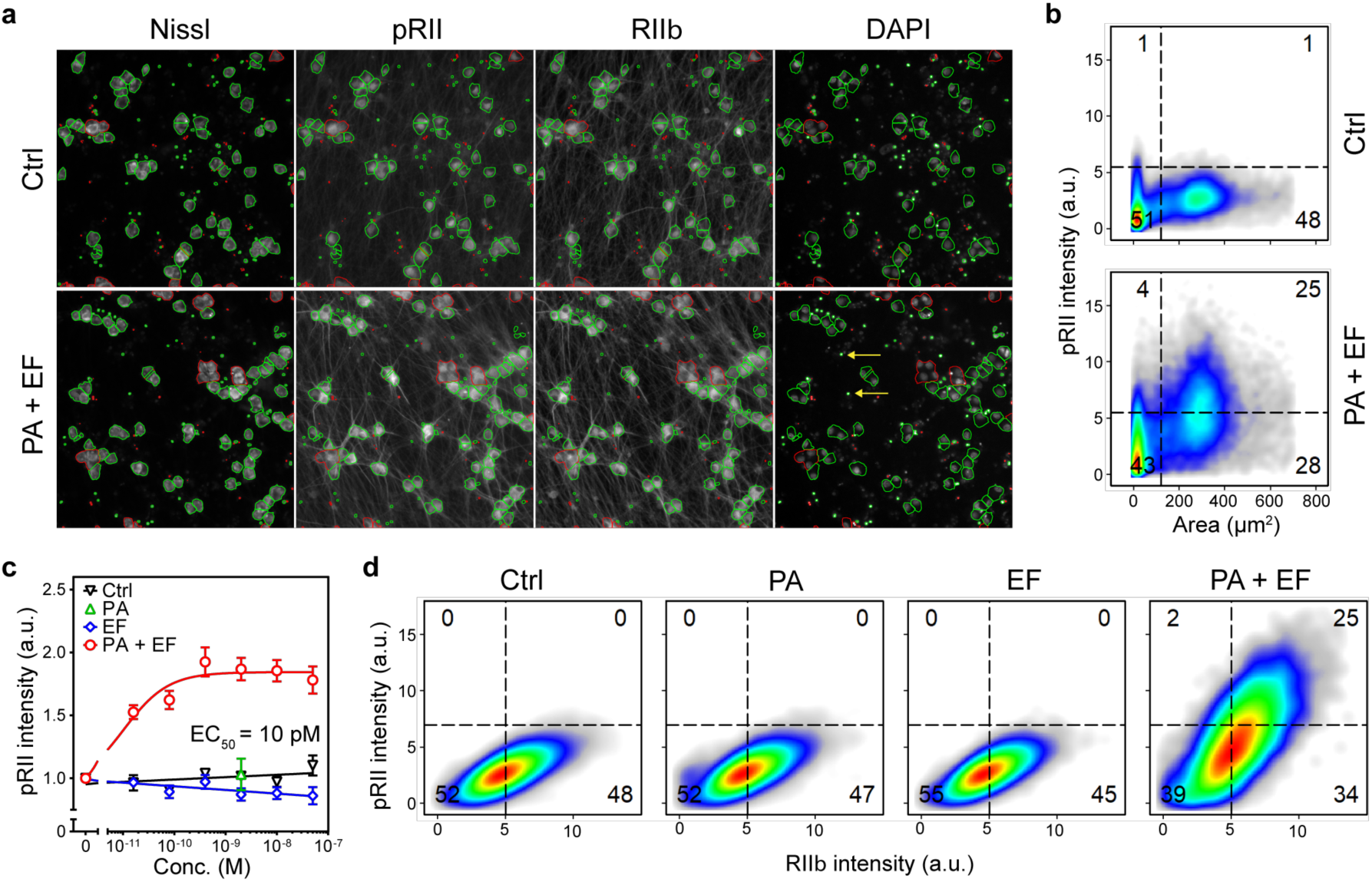
Edema Toxin acts on human iPSC-derived sensory neurons. **(a)** Representative HCS microscopy images of human iPSC-derived nociceptors stimulated with vehicle control (Ctrl) or EF + PA (10 nM each) for 2 h. Cultures were labeled with fluorescent Nissl to identify the cells, and pRII and RIIβ to quantify PKA-II signaling activity. Green or red encircled cells indicate automatically selected or rejected objects, respectively (see methods section). Scale bar, 200 µm. **(b)** Cultures of human iPS cells resulted in fully differentiated neurons with large cell bodies as well as smaller-sized specks (see yellow arrows in A), which were excluded from further analysis. **(c)** Dose-response curve of pRII intensity in human iPSC-derived nociceptors exposed to EF-A (0 - 50 nM, 2 h) in the absence or presence of a constant dose of PA (10 nM). Values represent means ± SEM; *n* = 8 wells from one culture of differentiated neurons analyzed in four replicate experiments; >8000 neurons/condition. **(d)** Single cell data of human iPSC-derived nociceptors stimulated with control solvent (0.1% BSA) or EF (2 nM) and PA (10 nM) for 2 h.

## DISCUSSION

Bacterial products can modulate nervous system function and may be utilized as molecular tools for targeting neuronal subtypes. Here, we describe a novel interaction between anthrax toxin and neurons based on nociceptor expression of the high-affinity receptor ANTXR2. Native ET and engineered anthrax toxins targeted nociceptive neurons for functional modulation, producing significant analgesia *in vivo* and demonstrating application as a molecular delivery platform. Our findings illustrate that identifying the novel ways bacterial molecules act on the nervous system is a rich area of research, both for basic understanding of bacterial pathogenesis and design of novel therapeutic strategies.

### Bacterial toxins modulate neuronal signaling and pain

A prototypic example of microbial toxins that have been utilized to target neurons in clinical applications are the botulinum neurotoxins produced by *Clostridium botulinum*, which inhibit neurotransmission and produce flaccid paralysis *(53)*. BoNT/A has now been formulated for use in cosmetic dermatology, as well as treatment of muscle spasticity and neurological conditions such as migraine. Sensory neurons in particular are poised to be targeted by bacterial molecules as they densely innervate barrier tissues. In addition to this work, several recent studies have illustrated how bacterial toxins can modulate pain. We have reported that *Staphylococcus aureus* and *Streptococcus pyogenes* pore-forming toxins induce ion influx and action potential firing in nociceptive neurons to produce pain during infection *(20–22)*. Other groups have identified analgesic mediators, such as a mycolactone isolated from *Mycobacterium ulcerans,* a pathogen that induces painless ulcers *(54)*, and a lipopeptide isolated from the gut probiotic *E. coli* strain Nissle *(55)*. Here we have identified anthrax toxins as novel modulators of nociceptive neurons and pain, and utilized their intoxication mechanism to deliver therapeutically relevant cargo into nociceptors.

### Physiological implications of ANTXR2 expression in nociceptive neurons

ANTXR2 is a single-pass transmembrane protein that is expressed broadly in the periphery, including various tissue types such as the heart, lung and liver *(11)*. The physiological role of ANTXR2 is not fully understood but may involve maintaining homeostasis of the extracellular matrix (ECM). ANTXR2 has been reported to bind to collagen type IV and laminin *(56)* and mediate the internalization and degradation of collagen type IV *(41)*. Global ANTXR2 knock-out mice develop normally but show parturition defects due to abnormal deposition of ECM proteins in the uterus *(57, 58)*. Loss-of-function mutations in ANTXR2 in humans cause Hyaline fibromatosis syndrome (HFS), characterized by the development of nodules containing excess ECM proteins *(59, 60)*.

In this study, we show that ANTXR2 is expressed in somatosensory neurons and enriched in nociceptive neurons. Although we found that conditional deletion of ANTXR2 from Nav1.8^+^ sensory neurons do not cause major defects in pain-like behavior in mice, the endogenous contributions of ANTXR2 to nociceptor function or development remain to be further investigated. To note, sensory or neurological abnormalities have not been reported in ANTXR2 knock-out mice or patients with HFS.

Beyond its physiological role, ANTXR2 is also a key mediator of anthrax pathogenesis. Expression of ANTXR2 on nociceptive neurons may render them targets of anthrax toxin during *B. anthracis* infection. Recent work has demonstrated that bacteria or bacterial toxins can directly stimulate nociceptive neurons during infection, triggering the release of neuropeptides such as CGRP that modulate pathogenesis *(21, 22, 61, 62)*. The initial sites of anthrax infection, including the skin, lung and gastrointestinal tract, are heavily innervated by nociceptors, placing them in proximity to locally produced toxins. Neurons at distal sites may also be targeted during toxemia when anthrax toxin enters circulation *(63, 64)*. It remains to be investigated whether nociceptors may play a potential role in anthrax pathogenesis by affecting neuronal signaling.

### Pain modulation by Edema Toxin and cAMP signaling

Here, we report that intrathecal administration of ET targets the DRG to elevate cAMP levels and activate PKA in neurons. The duration of these effects correlated with profound analgesia. Our observation that ET produces cAMP and analgesia concurrently contrasts with the traditional paradigm of inflammation-induced hyperalgesia. Prior work has shown that mediators such as prostaglandin E2 (PGE2) promote nociceptor sensitization by activating Gs proteins, endogenous adenylyl cyclases and PKA *(40)* to phosphorylate ion channels such as Trpv1 *(65)* and Nav1.8 *(33, 34)*. Accordingly, intrathecal or intra-ganglionic injections of membrane-permeable cAMP *(66, 67)* and intrathecal injection of forskolin *(68)* have been shown to produce mechanical hyperalgesia in rats, contrary to our observations with ET.

As a potent bacterial adenylyl cyclase, EF may induce qualitatively different cAMP responses than endogenous mediators. Endogenously generated cAMP is spatiotemporally restricted and compartmentalized in microdomains maintained by phosphodiesterases (PDEs) *(69)*. Downstream mediators of cAMP are also compartmentalized, such as PKA which is confined to subcellular locations by A-kinase anchoring proteins (AKAPs) *(70)*. EF is orders of magnitude more potent than mammalian adenylyl cyclases *(71)* and emanates cAMP from the perinuclear region where the toxin likely exits from late endosomes, creating a downhill gradient of cAMP towards the cell membrane *(72)*. While we observed at the cellular level that ET does activates PKA in nociceptive neurons in line with elevated cAMP, the exact nature of downstream pathways induced by large quantities of mislocalized cAMP remains to be examined. The involvement of other cAMP effectors such as EPAC, a guanine-nucleotide-exchange factor, also merits investigation.

Conditional deletion of ANTXR2 from Nav1.8-expressing nociceptors attenuated ET-induced analgesia, indicating the direct action of ET on nociceptors. However, we found that application of ET to nociceptive neurons in culture increased their excitability, consistent with PKA-mediated sensitization but contrary to the analgesic phenotype induced by intrathecal administration. Such a dichotomy has also been observed with a gain-of-function mutation in Nav1.9 which leads to loss of pain perception *(73)*. Potentially, our results may also support the involvement of secretory factors acting in an autocrine or paracrine manner. One possible mechanism could be secretion of cAMP into the extracellular space and subsequent conversion to adenosine *(74, 75)*, which can act on adenosine receptors on nociceptors to block pain *(76)*. Such a mechanism involving a diffusible mediator may explain the broad analgesic effects of ET across multiple models of pain, including those that may not exclusively involve C-fibers expressing ANTXR2. For instance, ET effectively silenced neuropathic pain induced by nerve injury, which has been suggested to be driven by Aβ mechanoreceptors that normally mediate innocuous touch *(77, 78)*. It is also possible that PKA activation could directly inhibit glutamate release at presynaptic terminals of nociceptors, similar to PKA inhibition of transmission seen at some hippocampal synapses *(79)*. In addition, we have not ruled out contributions from ET targeting other sensory neuron subtypes or non-neuronal cells via ANTXR2 or even ANTXR1.

In contrast to intrathecal administration of ET, intraplantar administration of the toxin produced mechanical hyperalgesia independent of ANTXR2 expression on neurons, accompanied by significant edema. ANTXR2 is expressed broadly in the periphery, including keratinocytes, smooth muscle cells, endothelial cells *(80)*, as well as innate and adaptive immune cells *(81)*, providing multiple targets for ET. ET has been reported to produce vascular leakage and edema through inducing inflammatory mediators such as histamine, prostanoids and substance P *(82)*, which are known to sensitize nociceptors *(83)* and may have mediated the hyperalgesia. The cellular source of such inflammatory mediators are yet unknown.

We additionally found that EF contributes to the development of mechanical allodynia during subcutaneous infection of the *B. anthracis* Sterne strain in mice. In humans, cutaneous anthrax is characterized by itchy blisters that make way to a painless eschar. By contrast, gastrointestinal anthrax produces ulcerative lesions and abdominal pain *(84)*. The modeling of infection in mice and mechanisms underlying these diverging pain phenotypes, including potential contributions from anthrax toxins, remain to be determined. However, it is clear that ET has major direct and indirect effects on pain *in vivo*.

The ability of ET to modulate pain presents broader implications for other bacterial toxins that also induce cAMP. Other adenylyl cyclase toxins include CyaA from *Bordetella pertussis (85)*, the etiologic agent of whooping cough, and ExoY from *Pseudomonas aeruginosa (86)*, an opportunistic pathogen that is a common cause of hospital-acquired infections. Other bacterial toxins act through G proteins to stimulate or disinhibit endogenous adenylyl cyclases, such as cholera toxin from *Vibrio cholera*; labile toxin from *Escherichia coli*; and pertussis toxin from *B. pertussis*. Whether such toxins can act on nociceptive neurons is unknown. Curiously, intrathecal administration of cholera toxin has been reported to attenuate inflammatory and neuropathic pain in rats *(87)*, raising the question of whether cAMP-inducing toxins can trigger a shared analgesic pathway.

### Potential analgesic strategies based on engineered anthrax toxins

Intrathecal ET produced broad and potent analgesia with minimal side effects, suggesting it may provide an avenue to treat pain with higher specificity compared to existing analgesics. For example, ziconotide, the synthetic version of ω-conotoxin, inhibits N-type calcium channels and is administered via intrathecal infusion devices to treat severe chronic pain. However, possibly due to channel expression in the brain, ziconotide can also cause psychiatric symptoms and neurological impairment, and is contraindicated in patients with preexisting history of psychiatric disorders *(88)*. ANTXR2 is nearly absent in the CNS, suggesting that the actions of ET may be more restricted to nociceptors compared to ziconotide. Such specificity may also reduce addictive potential compared to opioid-based painkillers, although this remains to be investigated.

Extending the duration of ET-induced analgesia may allow bolus administration and increased safety over infusion strategies. We observed that following ET injection, cAMP levels were elevated in relevant tissue for several hours before returning to baseline (**Fig. 3b, d**), suggesting that EF delivered to the cytoplasm becomes degraded or inactivated within hours. EF has been shown to be uniquely sensitive to ubiquitination *(89)* and susceptible to proteolytic degradation *(90)*. Although the exact intracellular half-life of EF is unknown, these observations suggest that the functional persistence of EF in the cytoplasm is fairly short. Methods that have successfully prolonged the intracellular half-life of LFN-DTA may potentially be applied to EF, such as attaching a D-amino acid at the N-terminus *(91)* or dimethylating all lysines *(92)* to avoid ubiquitination and proteosomal degradation. Other potential strategies may involve combination treatment with PDE inhibitors to prolong the duration of elevated cAMP.

Beyond native anthrax toxin, the PA + LFN system successfully delivered three separate enzymatic entities into nociceptive neurons for functional modulation, including the catalytic domains of diphtheria toxin, EF and botulinum toxin. Delivery of the botulinum toxin light chain produced significant blockade of neuropathic pain in the SNI model, illustrating the potential of the anthrax toxin system to target pain signaling. The flexibility of the PA pore and modular nature of the PA + LFN system make it an attractive delivery platform. A reported limitation of the PA + LFN system involves protein payloads that are very stable, which did not unfold efficiently to be translocated through the PA pore but could be successfully delivered with the introduction of destabilizing mutations *(93)*. In addition, the delivery capacity of the system remains to be fully characterized in nociceptive neurons, including the rate and extent of delivery. Our observation that repeated injections were required for PA + LFN-LC/A^C699S^ efficacy suggests that the delivery efficiency *in vivo* remains to be further optimized. Potential avenues to be explored include comparing the functionalities of the PA + LFN system and PA + EFN system, given the potent effects of ET that we observed *in vivo*. Judicial selection of protein cargoes functionally relevant in nociceptive neurons may add an additional layer of selectivity to this delivery strategy. Multiple LFN-linked cargo may be delivered simultaneously in a combinatorial manner for synergistic effects.

Altogether, we propose that the anthrax toxin system is uniquely positioned to manipulate nociceptive neurons and merits further development as a versatile intracellular delivery platform.

## MATERIALS AND METHODS

### Reagents

Forskolin (10 mM in DMSO) was purchased from Tocris (Bristol, UK). Lethal factor (LF) was purchased from List Biological Laboratories (#169, recombinant from *B. anthracis*). Protective antigen (PA) was obtained through BEI Resources, NIAID, NIH (#NR-140, recombinant from *B. anthracis*).

### Antibodies

The following antibodies were used in this study: Rabbit polyclonal anti-MEK-3 (1:500, Santa Cruz Biotechnology, #sc-961), rabbit polyclonal anti-p38 (1:1000, Cell Signaling Technology, #9212), rabbit monoclonal anti-phospho-p38 (Thr180/Tyr182) (clone D3F9, Cell Signaling Technology, #4511, 1:1000), rabbit polyclonal anti-SNAP-25 (1:1000, Millipore Sigma #S9684), rabbit polyclonal anti-GAPDH (1:30,000, Millipore Sigma #G9545), goat anti-rabbit IgG (1:1000, Cell Signaling Technology, #7074), chicken polyclonal anti-UCHL1 (1:2000, Novus, Cambridge, UK, #NB110-58872), rabbit monoclonal anti-RIIα (phospho-Ser96) (1:1000, clone 151, Abcam, Cambridge, UK, #ab32390), mouse monoclonal anti-RIIβ (1:2000, BD Transduction Laboratories, #610625), mouse monoclonal anti-NF200 (clone N52, Sigma, #N0142, 1:1000), mouse monoclonal anti-CaMKII alpha subunit (clone 6G9, Thermo Scientific, #MA1-048, 1:1000), mouse monoclonal anti-NaV1.8 (clone N134/12 Neuromab Facility, #75-166, 1:500), mouse monoclonal anti-CGRP (clone 4901, biorbyt, #orb319478, 1:500), goat polyclonal anti-TrkA, (R&D Systems, #AF1056, 1:500), goat polyclonal anti-TRPV1 (R&D Systems, #AF3066, 1:500), highly cross-adsorbed Alexa 647, 555, and 488 conjugated secondary antibodies (Invitrogen, Carlsbad, CA).

### Animals

All animal experiments were approved by the Institutional Animal Care and Use Committee (IACUC) at Harvard Medical School and were conducted in accordance with National Institutes of Health (NIH) animal research guidelines. C57BL/6J mice were purchased from Jackson Laboratory (Bar Harbor, ME) and bred at Harvard Medical School. Nav1.8-Cre mice were provided by John Wood (University College London) *(24)*. Antxr2^fl/fl^ mice in which the transmembrane domain of *Antxr2* is flanked by loxP sites were obtained from Jackson Laboratory (#027703), originally donated by Stephen Leppla (NIH) *(14)*. Mice lacking functional ANTXR2 in Nav1.8-lineage neurons were generated by crossing Nav1.8-Cre mice with Antxr2^fl/fl^ mice to obtain Nav1.8^cre/+^/Antxr2^fl/fl^ mice and control Nav1.8^+/+^/Antxr2^fl/fl^ littermates. Genotyping of this strain was performed as previously described *(9)*. Mice were kept on a 12 h light/dark cycle, provided *ad libitum* access to food and water and housed in individually ventilated microisolator cages within a full barrier, specific pathogen-free animal facility at Harvard Medical School. All experiments were performed with age- and sex-matched mice between 7 to 14 week of age unless otherwise noted. For high content screening (HCS) microscopy experiments, male C57Bl/6N mice (>24 g, aged 8-10 weeks) were obtained from Charles River. Mice were housed in a temperature- and humidity-controlled animal care facility at the University Hospital of Cologne on a 12 h light/dark cycle and provided with food and water *ad libitum*.

### Recombinant protein expression and purification

Sequences for all recombinant proteins are reported in **Supplementary Information**. The Edema Factor (EF) clone employed in this study contains an extra alanine at the N terminus compared to the native sequence, which has been shown to have a stabilizing effect on activity *(89)*. EF and LFN-DTA were expressed using the Champion pET SUMO expression system (ThermoFisher Scientific) in BL21(DE3) *E. coli* and purified using a HisTrap FF Ni-NTA column (GE Healthcare Life Sciences). The SUMO tag was cleaved by incubation with SUMO protease (Thermo Fisher Scientific) for 1 h at RT and removed by size-exclusion chromatography (SEC). EF underwent additional endotoxin removal by anion exchange chromatography (AEX) using a HiTrap Q HP anion exchange column (GE Healthcare Life Sciences). Endotoxin levels in the final product were measured using the Pierce LAL Chromogenic Endotoxin Quantitation Kit (ThermoFisher Scientific) to be 0.29 EU/mg.

LF_N_-EFC linking the N terminal domain of LF (residues 1 – 255 of native LF) and C terminal catalytic domain of EF (residues 258 – 766 of native EF) was cloned into the expression vector pSJ115 using standard molecular techniques. LFN-EFC and EF were expressed and purified from the avirulent *B. anthracis* strain BH460 as previously described *(90, 95)*.

For the LFN-LC/A^C699S^ construct, the N terminal domain of LF (residues 1 - 262 of native LF) and the catalytic domain of Botulinum neurotoxin type A1 (BoNT/A1) (residues 1 - 448 of native BoNT/A1) separated by a (GGS)2 linker was codon-optimized and cloned into the pK8 expression vector with a cleavable C-terminal His-tag. This was expressed in *E .coli* strain NiCo21 (DE3) using conditions previously described*(96)* and purified by IMAC using NiHP (GE Healthcare Life Sciences) and AEC using QHP (GE Healthcare Life Sciences) columns. The His-tag was removed by O/N incubation with TEV protease (Millipore Sigma) at 4 °C followed by negative INAC, and the final product was desalted into PBS pH 7.2 and stored at -80 °C.

### Analysis of published single-cell RNAseq, tissue expression and *in situ* hybridization data

Single-cell RNAseq data of mouse DRG neurons were obtained from Zeisel et al. *(25)* and Sharma et al. *(26)*. Average transcript levels of *Antxr2*, *Scn10a*, *Trpv1*, *Calca*, *Ntrk1*, *Ntrk2* and *Ntrk3* across DRG neuron clusters were obtained from mousebrain.org and plotted as a relative heat map using GraphPad Prism. The clusters were originally designated as PSPEP1-8, PSNF1-3 and PSNP1-6 by Zeisel et al. Expression of *Antxr2* and *Antxr1* in DRG neurons were plotted as a force-directed layout through https://kleintools.hms.harvard.edu/tools/springViewer_1_6_dev.html?datasets/Sharma2019/all and filtered for adult neurons only. Microarray data of *Antxr2* expression in the DRG and brain regions were obtained from BioGPS.org (http://biogps.org/#;goto=genereport&;id=71914) *(97, 98)*. *In situ* hybridization data of *Antxr2* in the adult brain (P56) and juvenile spinal cord (P4) were obtained from the Mouse Brain Atlas (http://mouse.brain-map.org/experiment/show/69526659) and Mouse Spinal Cord Atlas (http://mousespinal.brain-map.org/imageseries/detail/100019979.html) *(99)* from the Allen Institute.

### qPCR

For analysis of mouse tissue, animals were anesthetized with Avertin (Millipore Sigma) solution and perfused with PBS prior to harvest. RNA was isolated using the RNeasy mini kit (Qiagen). For analysis of human tissue, total brain or DRG RNA was obtained from Clontech (#636530 and #636150). Brain RNA was pooled from 4 Asian males, aged 21 – 29, with an unknown cause of death. DRG RNA was pooled from 21 Caucasian males and females, aged 16 – 65, who underwent sudden death. Per the supplier, RNA was isolated by a modified guanidium thiocyanate method and integrity and purity was confirmed using an Agilent 2100 Bioanalyzer. Reverse transcription was performed using the iScript cDNA Synthesis Kit (Bio-Rad). Quantitative real-time PCR was performed using the Power SYBR Green PCR Master Mix (ThermoFisher Scientific) on a StepOnePlus RT PCR system (Applied Biosystems) or a LightCycler 96 (Roche). Expression relative to *Gapdh* was calculated using the comparative CT method.

*mAntxr1* F: CTCGCCCATCAAGGGAAAACT

*mAntxr1* R: TACTTGGCTGGCTGACTGTTC

*mAntxr2* F: CAGTGAGCATTCAGCCAAGTTC

*mAntxr2* R: CTGCAATCCCATTGGTACATTCTG

Primers to measure *Antxr2* expression in Nav1.8/Antxr2 mice were designed to span the deleted region:

*mAntxr2* F (for conditional KO): ATTGCAGCCATCGTAGCTATTT

*mAntxr2* R (for conditional KO): GCCAAAACCACCACATCAAG

*mGapdh* F: GGGTGTGAACCACGAGAAATATG

*mGapdh* R: TGTGAGGGAGATGCTCAGTGTTG

*hAntxr2* F: TGTGTGGGGGAGGAATTTCAG

*hAntxr2* R: AGGATAGGTGCAGGACAAAGC

*hGapdh* F: TGGCATTGCCCTCAACGA

*hGapdh* R: TGTGAGGAGGGGAGATTCAGT

### *In situ* hybridization

For chromogenic detection of *Antxr2*, freshly dissected lumbar DRGs were post-fixed in 10% NBF (Fisher Scientific) for 2 hours at 4°C, dehydrated in 30% sucrose in PBS overnight at 4°C, embedded in OCT medium (Tissue-Tek) and frozen in a dry ice/isopentane bath. For fluorescent detection of *Antxr2*, *Scn10a*, *Pvalb* and *Tubb3*, DRGs were embedded and frozen immediately after dissection. All DRGs were cryo-sectioned to 12 μm and mounted onto Superfrost Plus slides (Fisher Scientific). *In situ* hybridization was performed using the RNAscope system (Advanced Cell Diagnostics) following manufacturer’s protocol. For chromogenic detection, sections were digested with Protease Plus for 5 min at RT and processed with the RNAscope 2.5HD RED detection kit using the probe Mm-Antxr2-C1 (#46851). For fluorescent detection, sections were digested with Protease IV for 30 min at RT and processed with the RNAscope Multiplex Fluorescent detection kit v2 using the following probe combinations: (a) Mm-Antxr2-C1 (#46851), Mm-Scn10a-C2 (#426011-C2) and Mm-Tubb3-C3 (#423391-C3) or (b) Mm-Antxr2-C1 (#46851), Mm-Tubb3-C2 (#423391-C2) and Mm-Pvalb-C3 (#421931-C3). Widefield images were acquired at 20x magnification on an Eclipse TE2000-E Inverted Fluorescence Microscope (Nikon). For quantification, cell boundaries were drawn manually in ImageJ based on the *Tubb3* signal. Each cell was scored as positive or negative for expression of *Antxr2*, *Scn10a* or *Pvalb* by a blinded observer. Results from 12 – 15 fields (4 – 5 fields from 3 separate mice) were summed for final analysis.

### Dorsal root ganglia neuron dissection and culture

Dorsal root ganglia cultures were prepared as previously described *(22)* with minor modifications. Briefly, adult mice 6 – 12 weeks of age were euthanized by CO2 asphyxiation and dorsal root ganglia (DRG) were harvested from all segments of the spinal cord. Following enzymatic dissociation in Collagenase A and Dispase II for 40 min at 37°C, DRGs were triturated with decreasing diameters of syringe needles (18G, 22G, 25G) and purified through a layer of 15% BSA in neurobasal media (NBM). The resulting pellet was filtered through a 70 μm strainer and resuspended in NBM supplemented with B27 (Thermo Fisher) and L-glutamine (Thermo Fisher). Cells were then seeded in laminin-coated tissue culture plates and cultured in the presence of 50 ng/μL nerve growth factor (NGF) (Thermo Fisher) unless otherwise noted.

### cAMP detection from DRG cultures

DRG neurons were prepared as described in **Dorsal root ganglia neuron dissection and culture** and seeded 4000 cells per well in 96 well plates. Following overnight culture, cells were treated with the indicated toxin components in NBM containing 50 ng/μL NBM for 2 h at 37 °C, lysed by manual scrapping in 0.1M HCl containing 0.5% triton X-100, and clarified by centrifugation. cAMP levels in clarified lysates were measured using the Direct cAMP ELISA kit from Enzo Life Sciences (#ADI-900-066) following manufacturer’s protocol. Absorbance was measured on a Synergy Mx multi-mode microplate reader (BioTek) and fitted using GraphPad Prism.

### cAMP detection from tissue

Mice were given intrathecal or intraplantar administration of PBS or ET (2 μg PA + 2 μg EF). At the indicated timepoint, animals were euthanized by CO2 asphyxiation and blood was drained by cardiac puncture. Lumbar DRGs, lumbar spinal cord or the glabrous skin of the footpad were harvested and homogenized in 0.1M HCl + 0.5% triton X-100 using glass beads and a TissueLyser II (Qiagen). Lysate was centrifuged at 18,000 xg and 4 °C for 40 – 60 min to pellet cell debris. cAMP levels in clarified lysates were measured using the Direct cAMP ELISA kit from Enzo Life Sciences (#ADI-900-066) following manufacturer’s protocol. The protein concentration in clarified lysates were measured using the Pierce BCA Protein Assay kit (Thermo Fisher Scientific) following manufacturer’s protocol. Absorbances were measured on a Synergy Mx multi-mode microplate reader (BioTek) and fitted using GraphPad Prism. The concentration of cAMP in each sample was normalized by corresponding protein concentration.

### Western Blot Analysis

DRG neurons were prepared as described in **Dorsal root ganglia neuron dissection and culture** and seeded 15,000 cells per well in 96 well plates. Following overnight culture, cells were treated with combinations of 10 nM PA, 10 nM LF and 10 nM EF in NBM containing 50 ng/μL for 24 h at 37 °C. Cells were then lysed by manual scrapping in cold NP40 lysis buffer (Thermo Fisher Scientific) containing Halt protease and phosphatase inhibitors (Thermo Fisher Scientific). Samples were separated by SDS-PAGE and transferred to a PVDF membrane, which was blocked in TBST + 5% non-fat milk for 1 h at RT and washed with TBST. The membrane was incubated with the following primary antibodies in TBST + 5% BSA at 4°C overnight: rabbit polyclonal anti-MEK-3 (1:500, Santa Cruz Biotechnology, #sc-961), rabbit polyclonal anti-p38 (1:1000, Cell Signaling Technology, #9212) or rabbit monoclonal anti-phospho-p38 (Thr180/Tyr182) (clone D3F9, Cell Signaling Technology, #4511, 1:1000). Incubation with HRP-linked goat anti-rabbit IgG (1:2000, Cell Signaling Technology, #7074) in TBST + 2.5% BSA was performed for 1 h at RT. Signal was developed using the SuperSignal West Pico chemiluminescent substrate (Thermo Fisher Scientific) and imaged on an Amersham Imager 600 (GE Healthcare Life Sciences). The membrane was then stripped with Restore western blot stripping buffer (Thermo Fisher Scientific) for 15 min at 37°C, and re-probed with rabbit polyclonal anti-GAPDH (1:30,000, Millipore Sigma #G9545) in TBST + 5% BSA. Band intensities were quantified using ImageJ and each target was normalized to its respective loading control.

### Protein synthesis assay

DRG neurons were prepared as described in **Dorsal root ganglia neuron dissection and culture** and seeded 15,000 cells per well in 96 well plates. Following overnight culture, cells were treated with 0.1 pM to 10 nM LFN-DTA, with or without 10 nM PA, in NBM containing 50 ng/μL NGF for 6 h at 37 °C. Cells were then washed with leucine-free F12K media and incubated with 20 μCi/mL ^3^H-leucine in the same media for 1 h at 37 °C. Cells were washed three times with PBS and lysed in MicroScint-20 (PerkinElmer). Scintillation counting was performed on a MicroBeta TriLux (PerkinElmer).

### Botulinum toxin light chain activity assay

Activity of botulinum toxin light chains were measured using the BoTest A/E BoNT detection kit (BioSentinel) following manufacturer’s instructions. Fluorescence was measured on a Synergy Mx multi-mode microplate reader (BioTek).

### CGRP release and SNAP-25 cleavage assays

DRG neurons were prepared as described in **Dorsal root ganglia neuron dissection and culture** and seeded 7500 cells per well in 96 well plates. Cells were cultured for a week in NBM + 50 ng/μL NGF and pulsed with an additional 10 mM cytosine arabinoside for three days during day 3 – 5 of culture. Cells were then incubated with 0.1 pM to 1 nM LFN-LC/A^C699S^, with or without 10 nM PA, for 24 h at 37°C. Cells were stimulated with Krebs-Ringer buffer containing 80 mM KCl for 10 min at 37°C. The concentration of CGRP in the supernatant was determined using a CGRP ELISA kit (Cayman Chemical) following manufacturer’s protocol. Remaining cells were lysed in Bolt LDS sample buffer (Thermo Fisher Scientific) containing 0.1M DTT (Thermo Fisher Scientific) and 0.125 units/μL Benzonase (Millipore Sigma). Samples were then separated by SDS-PAGE and transferred to a PVDF membrane, which was blocked with TBST + 5% non-fat milk (NFM) for 1 h at RT. The membrane was blotted with rabbit polyclonal anti-SNAP-25 (1:1000, Millipore Sigma #S9684) in TBST + 5% NFM and HRP-linked goat anti-rabbit IgG (1:1000, Cell Signaling Technology, #7074) in TBST + 5% NFM. Signal was developed using the SuperSignal West Pico chemiluminescent substrate (Thermo Fisher Scientific) and imaged on an Amersham Imager 600 (GE Healthcare Life Sciences). Band intensities were quantified using ImageJ and percent cleavage was calculated using the following formula: Cleaved SNAP-25 / (Uncleaved SNAP-25 + Cleaved SNAP-25).

### *B. anthracis* culture and infection

The *Bacillus anthracis* Sterne 7702 strain (BA663, #NR-9396) and its isogenic mutant lacking EF (BA695, #NR-9398) were obtained from ATCC. Spores for both strains were prepared as previously described *(94)* and stored at 4 °C. All procedures were approved by the Committee on Microbiological Safety (COMS) at Harvard Medical School and conducted under Biosafety Level 2 protocols and guidelines. For infection, *B. anthracis* spores were streaked on BHI agar plates and grown overnight at 37 °C. A day prior to infection, bacterial colonies were picked and inoculated into 2x SG medium and grown overnight at 37 °C, 250 rpm. The day of the experiment, the overnight culture was diluted 50-fold into fresh 2x SG medium and grown for 2 h until mid log phase was reached. The culture was then washed twice with PBS and stored in PBS on ice until injection. Mice were anesthetized with 3% isoflurane (Patterson Veterinary) with oxygen using a precision vaporizer, after which the indicated dose of bacteria was injected subcutaneously into the left hind paw in a 20 μL volume. The inoculum was plated after injection to confirm dosages.

### Pain behavioral tests

All behavioral tests were performed with age- and sex-matched animals by observers blinded to the treatment groups or genotypes. Treatment groups were randomized and evenly distributed across cages and sex.

#### Mechanical and heat hyperalgesia tests

Mechanical and thermal pain thresholds were measured as previously described *(22)* with minor modifications. Briefly, mice were habituated on the behavior apparatus for 2 consecutive days for 1 hour each. After habituation, 3 baseline measurements were obtained on separate days prior to treatment. Mechanical pain thresholds were measured using von Frey filaments and the up/down method *(100)*. Thermal pain thresholds were measured using the Hargreaves apparatus (Model 390G, IITC Life Science).

#### Formalin tests

Prior to formalin injection, mice were habituated for 45 min in individual chambers in an infrared Behavior Observation Box (iBOB) (Harvard Apparatus), which allows recording of mice in the dark independent of an observer. Mice were then given intraplantar injection of 10 μL of 5% neutral-buffered formalin (NBF) (Fisher Scientific) in 0.9% saline (Millipore Sigma). Injections were performed under light restraint without anesthesia. Mice were immediately placed in iBOB and recorded for 45 min. Videos were scored by a blinded observer to quantify the total time spent licking and shaking the injected paw in 5 min intervals. The first phase of the response was defined as 0 – 5 min and the second phase as 15 – 35 min post formalin injection.

#### Carrageenan model

Carrageenan (Millipore Sigma) was dissolved in 0.9% saline (Millipore Sigma) to a 2% concentration and autoclaved. Mice were given intraplantar injection of 20 μL of 2% carrageenan and mechanical pain thresholds were monitored as described above.

#### Spared Nerve Injury (SNI) model

Mice were anesthetized with 100 mg/kg ketamine (Patterson Veterinary) and 10 mg/kg xylazine (Patterson Veterinary) given via intraperitoneal injection. 1 mL of sterile saline (Patterson Veterinary) was injected subcutaneously between the shoulder blades, and petrolatum ophthalmic ointment (Patterson Veterinary) was applied. The surgical area was shaved and disinfected with betadine (Patterson Veterinary) and ethanol (VWR). The skin and muscle layer were opened and separated using blunt dissection to reveal the sciatic nerve. Under a stereomicroscope, the tibial and peroneal nerves were ligated with suture (Silk, 3-0, non-absorbable) and a short segment below the ligation was cut and removed. The muscle layer was closed and skin was sutured (Ethilon, 4-0, absorbable). Mice were placed on a heating pad to recover. Sham surgery was performed identically to the regular procedure except the sciatic nerve was exposed and the incision closed without carrying out the final lesion. Mechanical pain thresholds were monitored as described above.

### Rotarod

Motor function and coordination after intrathecal ET administration was measured using a Rotarod apparatus (Med Associates). Mice were habituated on 3 consecutive days at constant speed (5 rpm for 60 s per trial, 3 trials per day), and received intrathecal administration of PBS or ET (2 μg PA + 2 μg EF). At 2, 4 or 24 hours post-injection, mice were placed on an accelerating rotarod (4 to 40 rpm in 5 min) and the time until the first passive rotation or fall was recorded. Each measurement consisted of 3 trials which were performed 15 min apart and averaged.

### Mouse DRG recordings

#### Cell preparation

DRG neurons were prepared as described in **Dorsal root ganglia neuron dissection and culture** and cultured overnight in NBM containing 50 ng/μL NGF and 2 ng/μL GDNF. Cells were plated on laminin-coated glass coverslips and incubated at 37 °C (95% O2, 5% CO2) overnight. Cells were treated with 10 nM PA + 10 nM EF in NBM + 50 ng/μL NGF + 2 ng/μL GDNF or vehicle for 2–10 h at 37 °C. Recordings were made within 1 h after removal from treatment.

#### Whole cell current-clamp recordings

Small DRG neurons (membrane capacitance: 6.6 ± 0.5 pF) were recorded at room temperature. Membrane voltage was recorded using electrodes with resistances of 5-9 MOhm. To reduce pipette capacitance, the tips of electrodes were wrapped with Parafilm. Giga-ohm seals were formed in Tyrode’s solution containing 155 mM NaCl, 3.5 mM KCl, 1.5 mM CaCl2, 1 mM MgCl2, 10 mM HEPES, 10 mM glucose, pH 7.4 adjusted with NaOH. Internal solution contained 140 mM K aspartate, 13.5 mM NaCl, 1.6 mM MgCl2, 0.09 mM EGTA, 9 mM HEPES, 14 mM creatine phosphate (Tris salt), 4 mM MgATP, 0.3 mM Tris-GTP, pH 7.2 adjusted with KOH. Reported membrane potentials were corrected for a liquid junction potential of -10mV between the internal and extracellular solutions *(101)*. After breaking into the cells, pipette capacitance was compensated (50% - 60%) and bridge balance was set to compensate for series resistance. Membrane potential was recorded with an Axon Instruments Multiclamp 700B Amplifier (Molecular Devices) and voltages and currents were controlled and sampled using a Digidata 1322A interface using pClamp 9.2 software (Molecular Devices). Membrane voltage signals were filtered at 10 kHz and digitized at 100 kHz. Analysis was performed with Igor Pro (Wavemetrics, Lake Oswego, OR) using DataAccess (Bruxton Software) to import pClamp data. Collected data are presented as mean ± SEM.

### DRG neurons for HCS microcopy

Mice were euthanized between 9 – 12 AM by CO2 intoxication, and cervical, lumbar and thoracic DRGs were removed within 30 min per animal. DRGs were incubated in Neurobasal A/B27 medium (Invitrogen, Carlsbad, CA) containing collagenase P (Roche, Penzberg, DE) (0.2 U/mL, 1 h, 37°C, 5% CO2). DRGs were dissociated by trituration with fire-polished Pasteur pipettes. Axon stumps and disrupted cells were removed by bovine serum albumin (BSA) gradient centrifugation (15% BSA, 120 *g*, 8 min). Viable cells were resuspended in Neurobasal A/B27 medium, plated in 0.1 mg/mL poly-L-ornithine / 5 µg/mL laminin-precoated 96-well imaging plates (Greiner, Kremsmünster, AU) and incubated overnight (37°C, 5% CO2). Neuron density was 1500 neurons/cm^2^. DRG neurons were stimulated 24 h after isolation in 96-well imaging plates.

### Generation of iPSC-derived sensory neurons

Differentiation of iPSC into sensory neurons was performed as previously reported *(49)* with slight modifications. Briefly, single cell iPSCs were seeded at 3×10^5^ cells/cm^2^ in StemMAC iPS-Brew (Miltenyi Biotec, Bergisch Gladbach, DE) in presence of 10 µM Rock-Inhibitor Y-27632 (Cell Guidance Systems, Cambridge, UK) on Geltrex (Thermo Fisher Scientific, Waltham, USA) coated T175-flasks at day -1. After 24 h the medium was change to differentiation medium. Neural differentiation was initiated by dual-SMAD inhibition using 100 nM LDN 193189 (Axon Medchem BV, Groningen, NL) and 10 µM SB 431542 (Biozol, Eching, DE) from day 0 to day 6. To specify the differentiating cells into sensory neurons 3 µM CHIR 99021 (Miltenyi Biotec, Bergisch Gladbach, DE), 10 µM SU5402 (Sigma Aldrich, St. Louis, USA) and 10 µM DAPT (Axon Medchem BV, Groningen, NL) were added to the culture from day 3 to day 14. Two basal media were used during differentiation: Media1 consists of Knock out DMEM/F-12 (Thermo Fisher Scientific, Waltham, USA) with 20% Knock Out Serum Replacement (Thermo Fisher Scientific, Waltham, USA), 2 mM (1x) GlutaMAX (Thermo Fisher Scientific, Waltham, USA), 100 µM (1x) NEAA (Thermo Fisher Scientific, Waltham, USA) and 0.02 mM 2-Mercaptoethanol (Thermo Fisher Scientific, Waltham, USA). Media2 consists of Neurobasal Media (Thermo Fisher Scientific, Waltham, USA) supplemented with 1% N2 supplement (Thermo Fisher Scientific, Waltham, USA), 2% B27 supplement (Thermo Fisher Scientific, Waltham, USA), 2 mM (1x) GlutaMAX (Thermo Fisher Scientific, Waltham, USA) and 0.02 mM 2-Mercaptoethanol (Thermo Fisher Scientific, Waltham, USA). Between d0 and d3 cells were kept in Media1, from day 4 to 5 cells were kept in 75% Media1 and 25% Media2, for day 6 cells were kept in 50% Media1 and 50% Media2, from day 7 to 9 cells were kept in 25% Media1 and 75% Media2 and from day 10 to 14 differentiating cells were kept in 100% Media2. At day 14 of differentiated cells were dissociated with Accutase (Thermo Fisher Scientific, Waltham, USA) and frozen in cold CryoStor CS10 freezing medium (Sigma Aldrich, St. Louis, USA) at -80 °C. After 24 h frozen cells were transferred to a liquid nitrogen tank for long-term storage.

Cells were plated after thawing in Geltrex coated 96-well imaging plates at a density of 40.000 cells per well in maturation medium. Maturation medium consisting of Neurobasal A (Thermo Fisher Scientific, Waltham, USA) supplemented with 1% N2 supplement (Thermo Fisher Scientific, Waltham, USA), 2% B27 supplement (Thermo Fisher Scientific, Waltham, USA), 2 mM (1x) GlutaMAX (Thermo Fisher Scientific, Waltham, USA), 0.02 mM 2-Mercaptoethanol (Thermo Fisher Scientific, Waltham, USA), 12 µg/mL Gentamycin (Thermo Fisher Scientific, Waltham, USA), 200 µM Ascorbic acid (Sigma Aldrich, St. Louis, USA), 0.1 µg/mL human recombinant Laminin LN521 (BioLamina, Sundyberg, SWE), 10 ng/mL GDNF (Cell Guidance Systems, Cambridge, UK), 10 ng/mL BDNF (Cell Guidance Systems, Cambridge, UK), 10 ng/mL NGF (Peprotech, Hamburg, DE) and 10 ng/mL NT3 (Peprotech, Hamburg, DE). Four days after plating cultures were treated with 1µg/mL Mitomyin C (Sigma Aldrich, St. Louis, USA) for 2 h at 37 °C to inactivate proliferative cells. Medium was changes twice per week. Cells were matured for three weeks prior to stimulation.

### Cell culture stimulation and fixation for HCS microscopy

DRG neurons and human IPSC-derived sensory neurons were stimulated in 96-well imaging plates. Compounds were dissolved in 12.5 μL PBS in 96-well V-bottom plates, mixed with 50 μL medium from the culture wells, and added back to the same wells. Stimulations were performed with automated 8 channel pipettes (Eppendorf, Hamburg, DE) at low dispense speed on heated blocks, and stimulated cells were placed back in the incubator. The cells were fixed for 10 minutes at RT by adding 100 μL of 8% paraformaldehyde resulting in a final concentration of 4%.

### Quantitative HCS microscopy

Fixed DRG cells were treated with goat or donkey serum blocking (2% serum, 1% BSA, 0.1% Triton X-100, 0.05% Tween 20, 1 h, RT) and incubated with respective primary antibodies diluted in 1% BSA in PBS at 4 °C overnight. After three washes with PBS (30 min, RT), cells were incubated with secondary Alexa dye-coupled antibodies (1:1000, 1 h, RT). After three final washes (30 min, RT), wells of 96-well plates were filled with PBS, sealed, and stored at 4 °C until scanning. For IPSC-derived cultures the Nissl staining was performed after primary and secondary staining. Wells were rinse with 0.1% Triton X-100 (5 min, RT) in PBS followed by 20 min incubation with Nissl solution (1:500, 20 min, RT). Three rinsing steps were performed as follow: 5 min PBS, 5 min 0.1% Triton X-100, 30 min three final washes with PBS. Latest, wells were filled with PBS, sealed, and stored at 4 °C.

We used a Cellomics ArrayScan XTI microscope equipped with an X1 CCD camera and LED light source to scan stained cultures of DRG neurons in 96-well imaging plates. 2×2 binned images (1104 x 1104 pixels) were acquired with 10× (NA = 0.3) EC Plan Neo-Fluor objective (Zeiss) for DRG and 20x objective for IPSC-derived nociceptors. Images were analyzed using the Cellomics software package. Briefly, images of UCHL1/Nissl staining were background corrected (low pass filtration), converted to binary image masks (fixed threshold), segmented (geometric method), and neurons were identified by the object selection parameters: size of 80-7500 µm^2^ (DRG), 10–600 µm^2^ (IPSC-SN), circularity (perimeter^2^ / 4π area) of 1-3; length-to-width ratio of 1-2 (DRG), 1-3 (IPSC-SN); average intensity of 800-12000 (DRG), 300-5000 (IPSC-SN), and total intensity of 2×10^5^-5x10^7^ (DRG), 500-10^9^ (IPSC-SN). The image masks were then used to quantify signal in other channels. To calculate spill-over between fluorescence channels, three respective controls were prepared for each triple staining: (i) UCHL1/Nissl alone, (ii) UCHL1/Nissl + antibody 1, and (iii) UCHL1/Nissl + antibody 2. Raw fluorescence data of the controls were used to calculate the slope of best fit straight lines by linear regression, which was then used to compensate spill-over as described previously *(102)*. Compensated data were scaled to a mean value of 1 for the unstimulated cells to adjust for variability between experimental days. One and two-dimensional probability density plots were generated using R packages. Gating of subpopulations was performed by setting thresholds at local minima of probability density plots. Statistical analyses were performed with Students t-tests, one-, or two-way ANOVA with respective post hoc tests as indicated in the figure legends. P < 0.05 was considered statistically significant. Dose-response curves from HCS microscopy were generated using non-linear regression curve-fitting (three parameter, standard Hill slope) using Prism software (GraphPad, La Jolla, CA). The parameters of the model (top, bottom, or pEC50/pIC50 values) were compared using the extra-sum-of-squares F test. High content screening kinetic experiments were analyzed with R using ordinary two-way ANOVA. Bonferroni’s post hoc analysis was applied to determine p values of selected pairs defined in a contrast matrix using the R library multcomp. Error bars represent the standard error of the mean (SEM) of three independent replicate experiments using cells of different animals.

### Analysis of spinal cord and DRG sections for PKA activation

Mice received intrathecal injection of PBS or ET (2 μg PA + 2 μg EF). At 2 h post-injection, L3 – L6 DRGs or lumbar spinal cord were harvested and fixed with 4% PFA (Electron Microscopy Sciences) in PBS for 2 h on ice and dehydrated in 30% sucrose in PBS at 4 °C overnight. Lumbar spinal cord was fixed with 4% PFA (Electron Microscopy Sciences) in PBS overnight and dehydrated in 30% sucrose in PBS at 4 °C for 2 – 3 days. Both tissue were embedded in OCT medium (Tissue-Tek) and snap frozen in an isopentane/dry ice bath. Frozen blocks were cut into 12 µm sections, mounted on slides, dried for 30 minutes at RT, and stored at -80 °C. Thawed sections were post-fixed in 2% PFA for 10 minutes at 4 °C, rinsed in PBS for 30 minutes, incubated in goat serum blocking (2% serum, 1% BSA, 0.1% Triton X-100, 0.05% Tween 20, 1h, RT), and incubated with respective primary antibodies diluted in 1% BSA in PBS at 4 °C overnight. After three washes with PBS (30 min, RT), sections were incubated with secondary Alexa dye-coupled antibodies (1:1000, 1h, RT). After three final washes (30 min, RT), the sections were mounted with Fluoromount-G (Southern Biotech) containing DAPI (0.5 µg/mL).

Images of sections were acquired with a Leica DM6000B epifluorescence microscope controlled by Leica MetaMorph (version 1.4.0) using a 20x objective (Leica HC PLAN APO, 0.7). The images were analyzed using a custom Fiji Image J plugin. Briefly, UCHL1 images (8 bit) were background corrected (rolling ball radius of 500 pixel), blurred (Gaussian with 1 pixel), and contrast enhanced (Enhance contrast value of 0.7). Then a fixed threshold was applied (40-250) followed by watershedding and particle analysis (size of 1000-20000, circularity of 0.3-1). The obtained image mask was transferred to the background corrected pRII images. To quantify pRII signal intensities in images of spinal cord sections, RIIβ-positive regions of dorsal horns were selected and used as regions of interest to quantify pRII intensities.

### Statistical analysis

All statistical analyses were performed using GraphPad Prism software. Data are represented as mean ± standard error (SEM) unless otherwise noted. Sample size was chosen based on standard practice in the field and are reported in figure legends. Statistical tests are reported in figure legends.

## SUPPLEMENTARY MATERIALS

**Fig. S1:**
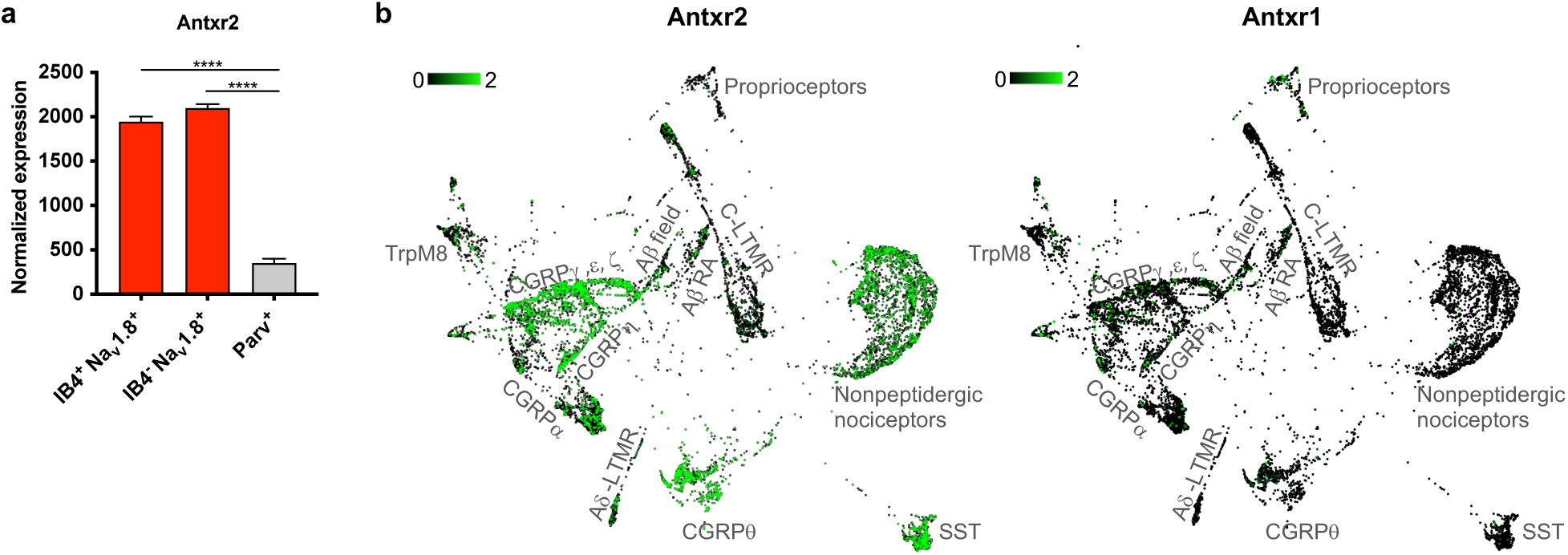
A*n*txr2 but not *Antxr1* is enriched in nociceptive neurons in the DRG. **(a)** Microarray analysis of *Antxr2* expression in sorted DRG neuron subsets *(1)*. **(b)** Force-directed layout of adult DRG sensory neurons overlaid with expression levels of *Antxr2* or *Antxr1,* from single cell RNA-seq data *(2)*. SST, somatostatin.

**Fig. S2:**
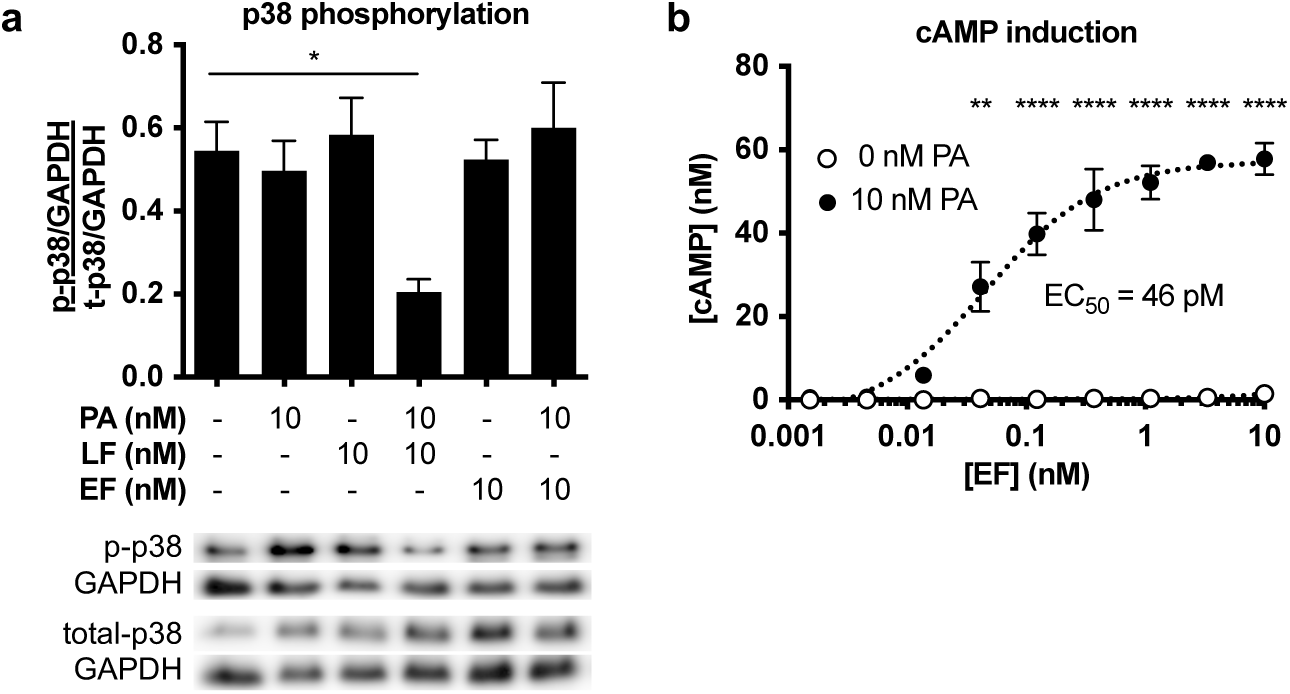
Anthrax toxin alters intracellular signaling in DRG neurons. **(a)** Phosphorylation levels of p38 in DRG cultures treated with PA, LF or EF for 24 h. Quantifications are from four independent experiments. **(b)** cAMP levels in DRG cultures treated with PA, LF or EF for 2 h. Results are from three independent experiments. **Statistical analysis: (a)** One-way ANOVA with Tukey’s post-test. **(b)** Two-way ANOVA with Sidak’s post-test. \**p* < 0.05, \*\**p* < 0.01, \*\*\*\**p* < 0.0001.

**Fig. S3:**
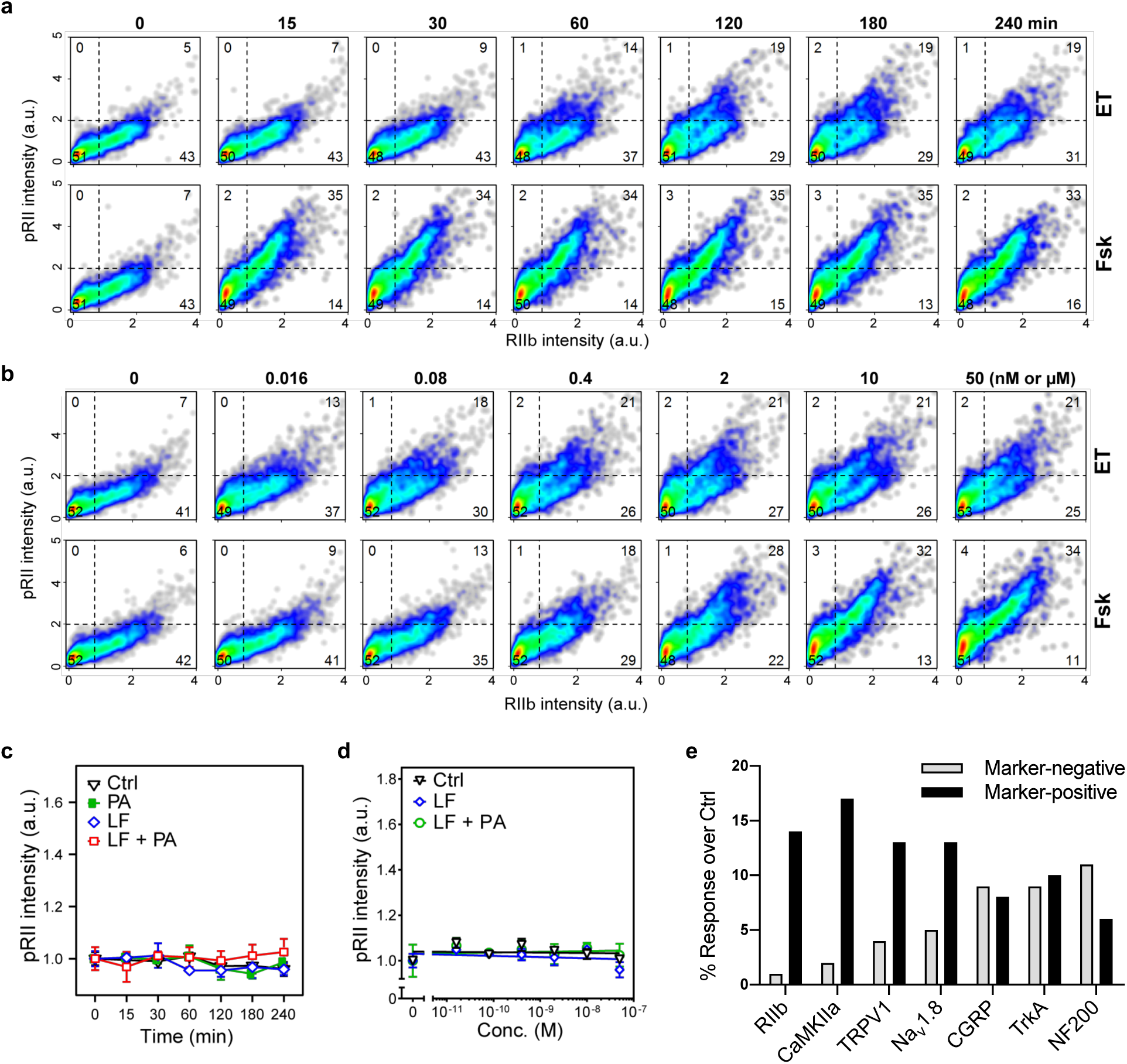
Edema Toxin induces PKA activity in nociceptive neurons. **(a)** Single cell data of DRG neurons stimulated with ET (10 nM EF + 10 nM PA) or Fsk (10 µM) for the indicated time points. Mean values are shown in Fig. 2e and g. **(b)** Single cell data of DRG neurons exposed to increasing doses of EF (0 - 50 nM) in the presence of PA (10 nM) or Fsk (0 - 50 µM) for 2 h. Mean values are shown in Fig. 2f and h. Combined data of n=3 replicate experiments are shown, >2500 neurons/condition in total. **(c)** Time-course of pRII intensity in DRG neurons stimulated with Ctrl (0.1% BSA), PA (10 nM), LF (10 nM) or the combination of both factors. **(d)** Dose-response curve of pRII intensity in DRG neurons exposed to LF (0 - 50 nM, 2 h) in the presence of a constant dose of PA (10 nM). **(e)** Changes in the percentage of pRII positive cells with ET treatment over Ctrl in each neuronal subgroup. The % response with Ctrl treatment was subtracted from the % response with ET treatment. Cells were treated with 10 nM PA + 10 nM EF for 2 h. Corresponding single cell data is shown in Fig. 2i.

**Fig. S4:**
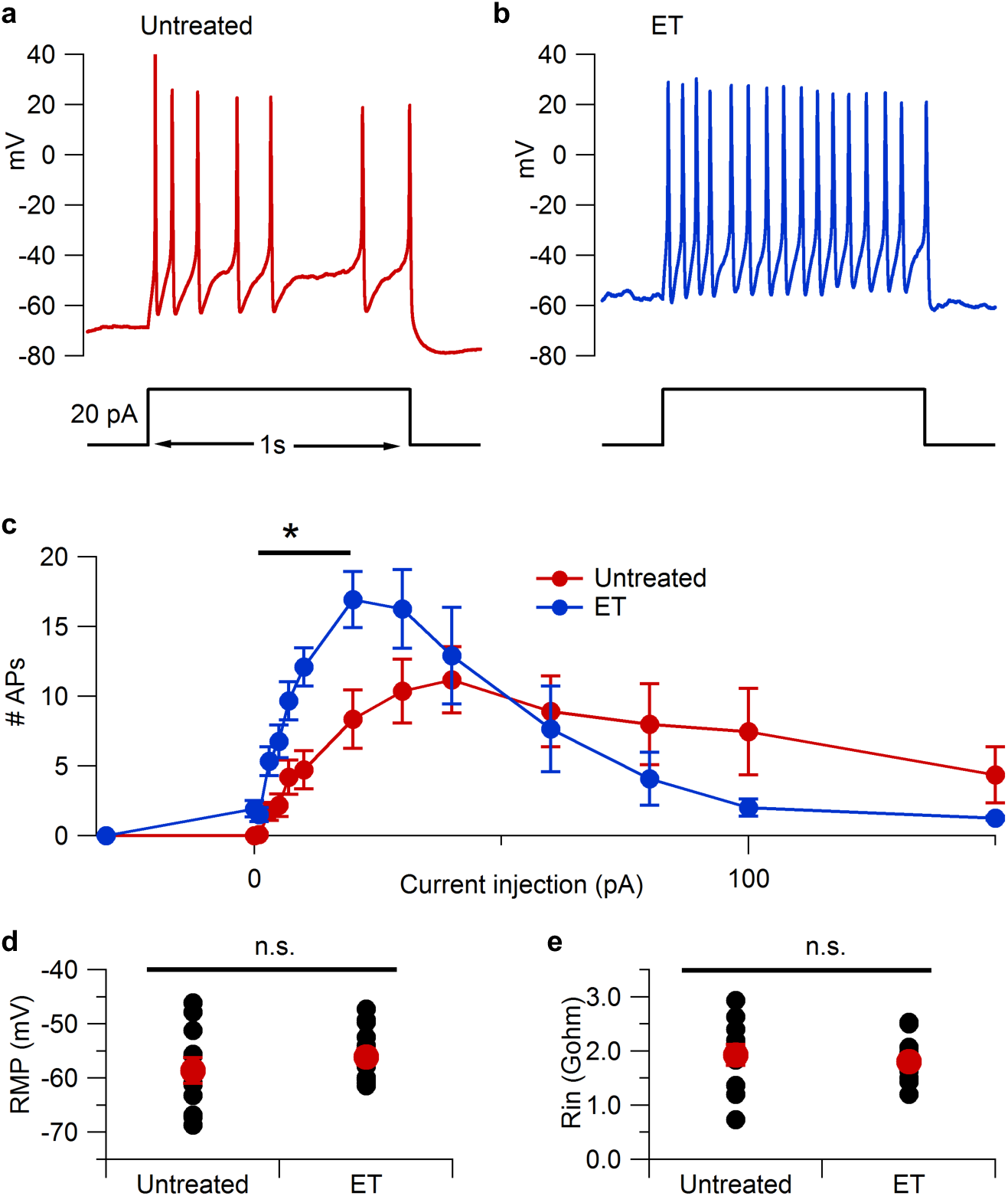
Edema Toxin treatment enhances excitability of small-diameter mouse DRG neurons. Excitability of small diameter DRG neurons was examined after treatment with control media (Untreated) or ET (10 nM PA + 10 nM EF) for 2 – 10 h at 37°C. **(a)** Action potential firing elicited by a 1-s 20-pA current injection in a control (vehicle-treated) small DRG neuron. **(b)** Action potential firing elicited by the same current injection in a representative ET-treated neuron. **(c)** Number of action potentials during 1-s current injections as a function of injected current (untreated group: n = 11; ET group: n = 12). Data plotted as mean ± SEM. **(d)** Resting potential of untreated (n=11) and ET-treated (n=12) neurons. **(e)** Input resistance of untreated (n=11) and ET-treated (n=12) neurons. **Statistical analysis: (c)** Two-way ANOVA with Sidak’s post-test, \**p* < 0.05. **(d, e)** Unpaired t-test, n.s, not significant.

**Fig. S5:**
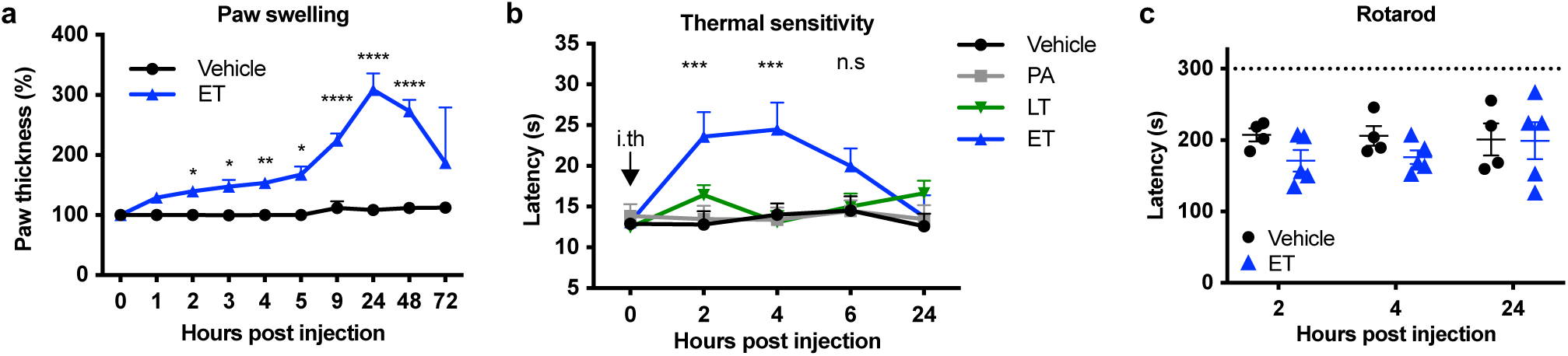
Edema Toxin modulates edema and pain *in vivo*. **(a)** Paw thickness after intraplantar administration of vehicle (PBS) or ET (2 μg PA + 2 μg EF) (n = 6/group). **(b)** Thermal sensitivity after intrathecal administration of vehicle (PBS), PA (2 μg), LT (2 μg PA + 2 μg LF) or ET (2 μg PA + 2 μg EF) (n = 8/group). **(c)** Motor function and coordination assessed by the rotarod test after intrathecal administration of vehicle (PBS) or ET (2 μg PA + 2 μg EF) (n = 4-5/group). **Statistical analysis: (a, c)** Two-way ANOVA with Sidak’s post-test. **(b)** Two-way ANOVA with Dunnett’s post-test, Vehicle vs. ET. \**p* < 0.05, \*\**p* < 0.01, \*\*\**p* < 0.001, \*\*\*\**p* < 0.0001, n.s, not significant.

**Fig. S6:**
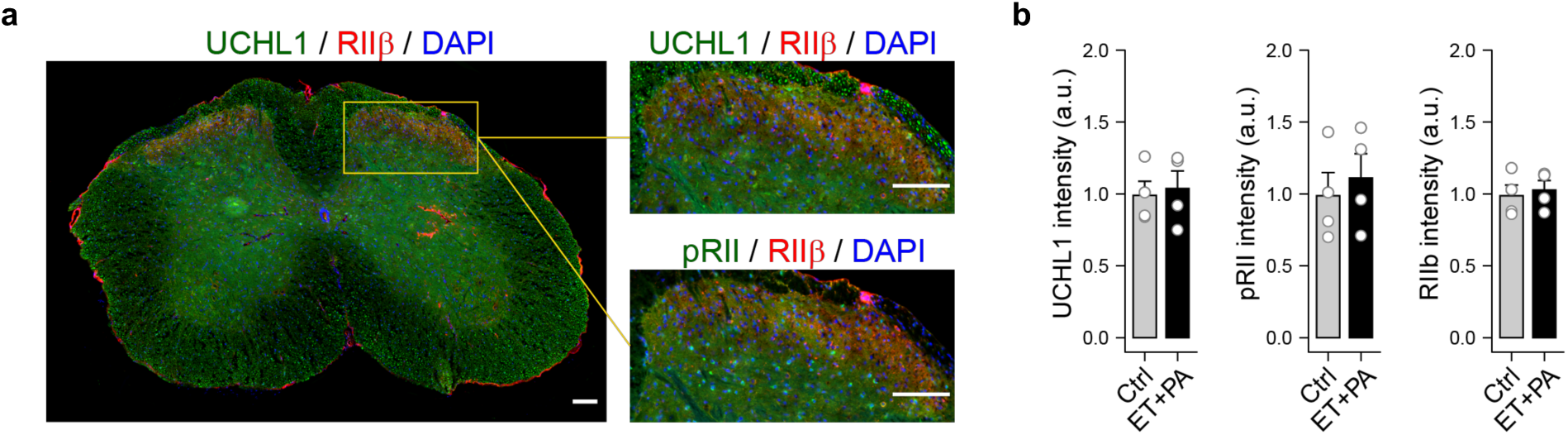
Edema Toxin does not induce detectable PKA activity in the spinal cord. **(a)** Representative spinal cord section obtained from mice 2 h post intrathecal injection of control solvent or EF-A and PA. RIIβ labels the nociceptive terminals in the outer layers of the dorsal horn. For quantification, RIIβ-positive dorsal horn areas were selected in merged images (see enlarged image sections). **(b)** Mean UCHL1, pRII and RIIβ intensity values of the analyzed spinal cord sections (n = 4 animals per group, 20 ± 3 dorsal horn areas per animal).

**Fig. S7:**
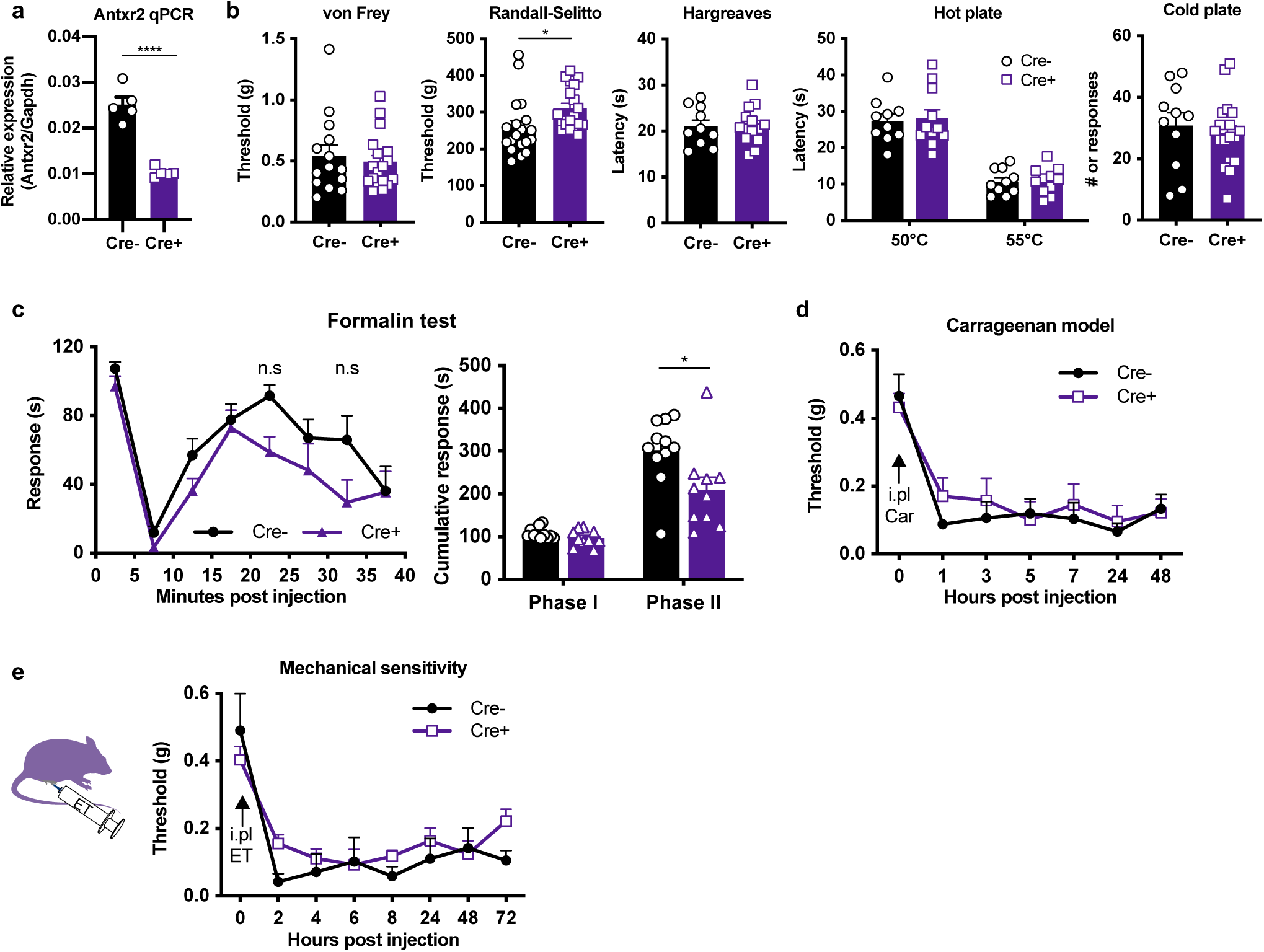
Neuronal ablation of ANTXR2 does not significantly affect pain behavior at baseline or following intraplantar administration of Edema Toxin. **(a)** qPCR analysis of Antxr2 expression in DRGs harvested from Nav1.8^cre/+^/Antxr2^fl/fl^ or Nav1.8^+/+^/Antxr2^fl/fl^ mice (n = 4-5/group). **(b)** Baseline pain behaviors compared between Nav1.8^cre/+^/Antxr2^fl/fl^ mice and Nav1.8^+/+^/Antxr2^fl/fl^ littermates (n = 10-19/group). **(c) (Left)** Acute pain behavior measured in 5 min intervals in Nav1.8^cre/+^/Antxr2^fl/fl^ or Nav1.8^+/+^/Antxr2^fl/fl^ mice following intraplantar injection of 5% formalin. **(Right)** Cumulative responses during Phase I (0 – 5 min) or Phase II (15 – 35 min) of formalin-induced pain (n = 10-11/group). **(d)** Mechanical sensitivity measured with von Frey filaments in Nav1.8^cre/+^/Antxr2^fl/fl^ or Nav1.8^+/+^/Antxr2^fl/fl^ mice following intraplantar injection of 20 μL of 2% carrageenan in saline (n = 5-6/group). **(e)** Mechanical sensitivity after intraplantar administration of ET (2 μg PA + 2 μg EF) to Nav1.8^cre/+^/Antxr2^fl/fl^ or Nav1.8^+/+^/Antxr2^fl/fl^ mice (n = 5-7/group). **Statistical analysis: (a, b)** Unpaired t-test. (**c, left)** Two-way ANOVA with Sidak’s post-test. **(c, right)** Unpaired t-test. **(d, e)** Two-way ANOVA with Sidak’s post-test. \**p* < 0.05, \*\*\*\**p* < 0.0001.

**Fig. S8:**
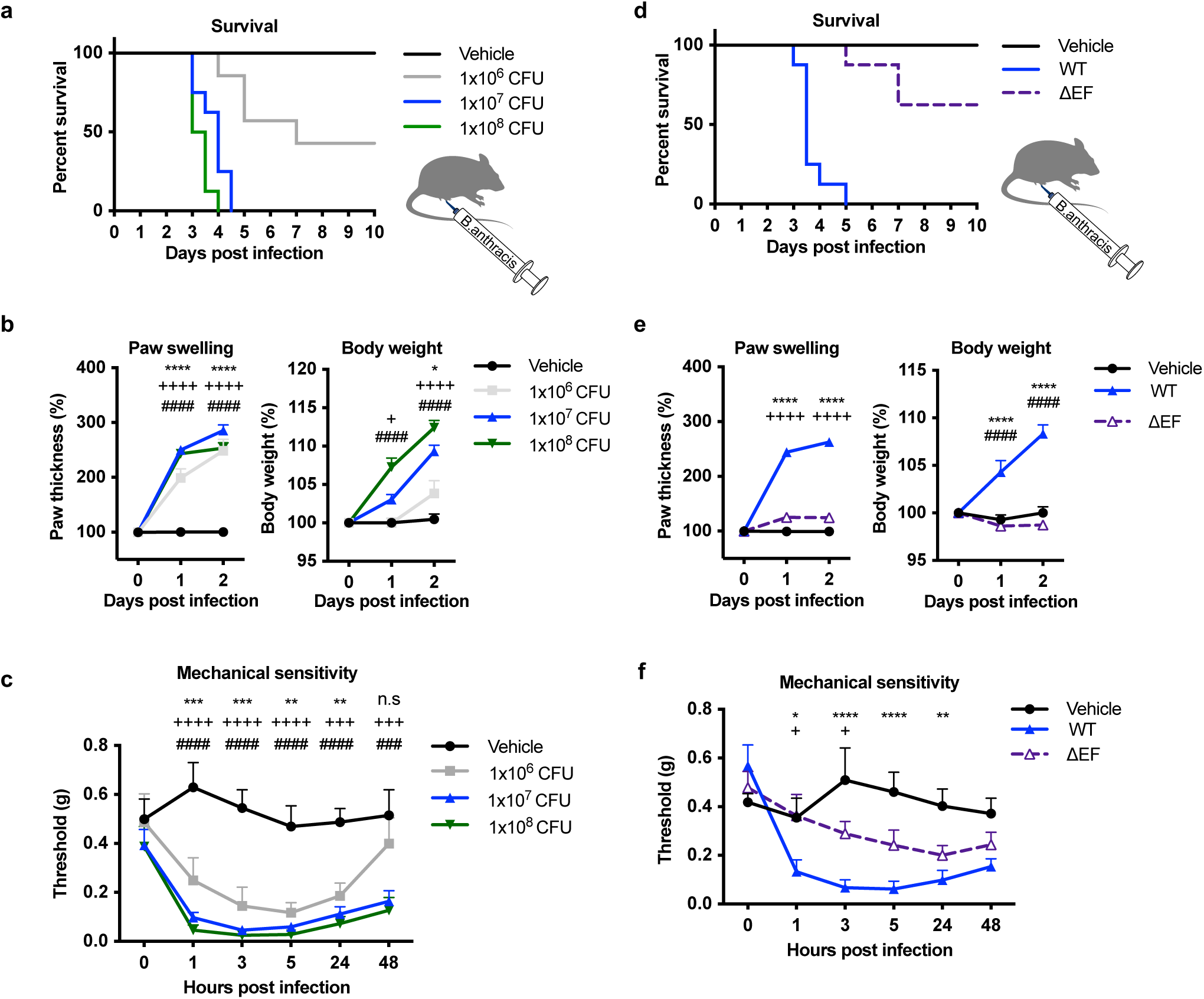
Edema Factor contributes to mechanical allodynia and inflammation during subcutaneous *B. anthracis* infection. **(a-c)** Vegetative wild-type *B. anthracis* Sterne 7702 at the indicated inoculums were injected subcutaneously into the footpad of mice. **(a)** Survival, **(b)** paw swelling and body weight changes post infection. **(c)** Mechanical sensitivity post infection measured with von Frey filaments (n = 7-8/group). **(d-f)** Vegetative wild-type *B. anthracis* Sterne 7702 (WT) or an isogenic mutant lacking EF (*Δ* EF) were injected subcutaneously into the footpad of mice at an inoculum of 1 *ι* × 10^7^ CFUs. **(d)** Survival, **(e)** paw swelling and body weight changes post infection. **(f)** Mechanical sensitivity post infection measured with von Frey filaments (n = 8/group). **Statistical analysis: (a, d)** Log-rank (Mantel-Cox) test. **(b, c)** Two-way ANOVA with Dunnett’s post-test. Vehicle vs. 1 × 10^6^ CFU, \**p* < 0.05, \*\**p* < 0.01, \*\*\**p* < 0.001, \*\*\*\**p* < 0.0001; Vehicle vs. 1 × 10^7^ CFU, +*p* < 0.05, +++*p* < 0.001, ++++*p* < 0.0001; Vehicle vs. 1 × ^8^ CFU, ###*p* < 0.001, ####*p* < 0.0001. **(e, f)** Two-way ANOVA with Tukey’s post-test. Vehicle vs. WT, \**p* < 0.05, \*\**p* < 0.01, \*\*\*\**p* < 0.0001; Vehicle vs. *Δ* EF, ++++*p* < 0.0001; WT vs. *Δ* EF, #*p* < 0.05, ####*p* < 0.0001.

**Fig. S9:**
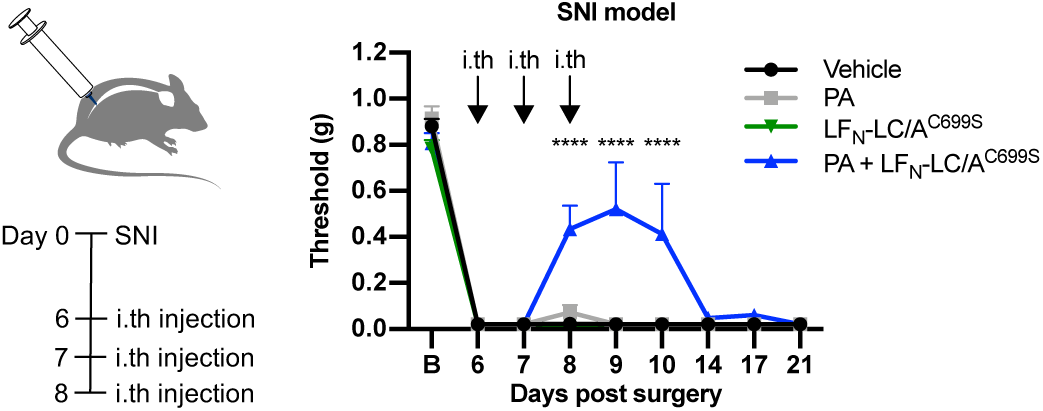
Anthrax toxin mediated delivery of botulinum toxin light chain blocks SNI-induced neuropathic pain. Mechanical sensitivity in SNI mice following intrathecal administration of vehicle (Gelatin Phosphate Buffer), PA (500 ng), LFN-LC/A^C699S^ (200 ng) or PA + LFN-LC/A^C699S^ (500 ng + 200 ng, respectively) for 3 consecutive days (n = 5-6/group). **Statistical analysis:** Two-way ANOVA with Tukey’s post-test. Vehicle vs. PA + LFN-LC/A^C699S^, \*\*\*\**p* < 0.0001.

**Fig S10. Protein sequences**

**EF** ANEHYTESDIKRNHKTEKNKTEKEKFKDSINNLVKTEFTNETLDKIQQTQDLLKKIPKDVLEIYSELGGEIYF TDIDLVEHKELQDLSEEEKNSMNSRGEKVPFASRFVFEKKRETPKLIINIKDYAINSEQSKEVYYEIGKGISLD IISKDKSLDPEFLNLIKSLSDDSDSSDLLFSQKFKEKLELNNKSIDINFIKENLTEFQHAFSLAFSYYFAPDHRT VLELYAPDMFEYMNKLEKGGFEKISESLKKEGVEKDRIDVLKGEKALKASGLVPEHADAFKKIARELNTYI LFRPVNKLATNLIKSGVATKGLNVHGKSSDWGPVAGYIPFDQDLSKKHGQQLAVEKGNLENKKSITEHEG EIGKIPLKLDHLRIEELKENGIILKGKKEIDNGKKYYLLESNNQVYEFRISDENNEVQYKTKEGKITVLGEKF NWRNIEVMAKNVEGVLKPLTADYDLFALAPSLTEIKKQIPQKEWDKVVNTPNSLEKQKGVTNLLIKYGIER KPDSTKGTLSNWQKQMLDRLNEAVKYTGYTGGDVVNHGTEQDNEEFPEKDNEIFIINPEGEFILTKNWEM TGRFIEKNITGKDYLYYFNRSYNKIAPGNKAYIEWTDPITKAKINTIPTSAEFIKNLSSIRRSSNVGVYKDSGD KDEFAKKESVKKIAGYLSDYYNSANHIFSQEKKRKISIFRGIQAYNEIENVLKSKQIAPEYKNYFQYLKERIT NQVQLLLTHQKSNIEFKLLYKQLNFTENETDNFEVFQKIIDEK

**LF_N_-DTA** AGGHGDVGMHVKEKEKNKDENKRKDEERNKTQEEHLKEIMKHIVKIEVKGEEAVKKEAAEKLLEKVPSD VLEMYKAIGGKIYIVDGDITKHISLEALSEDKKKIKDIYGKDALLHEHYVYAKEGYEPVLVIQSSEDYVENT EKALNVYYEIGKILSRDILSKINQPYQKFLDVLNTIKNASDSDGQDLLFTNQLKEHPTDFSVEFLEQNSNEVQ EVFAKAFAYYIEPQHRDVLQLYAPEAFNYMDKFNEQEINLSLEELKDQRSGRELERGADDVVDSSKSFVM ENFSSYHGTKPGYVDSIQKGIQKPKSGTQGNYDDDWKGFYSTDNKYDAAGYSVDNENPLSGKAGGVVKV TYPGLTKVLALKVDNAETIKKELGLSLTEPLMEQVGTEEFIKRFGDGASRVVLSLPFAEGSSSVEYINNWEQ AKALSVELEINFETRGKRGQDAMYEYMAQASAGNR

**LF_N_-EF_C_** AGGHGDVGMHVKEKEKNKDENKRKDEERNKTQEEHLKEIMKHIVKIEVKGEEAVKKEAAEKLLEKVPSD VLEMYKAIGGKIYIVDGDITKHISLEALSEDKKKIKDIYGKDALLHEHYVYAKEGYEPVLVIQSSEDYVENT EKALNVYYEIGKILSRDILSKINQPYQKFLDVLNTIKNASDSDGQDLLFTNQLKEHPTDFSVEFLEQNSNEVQ EVFAKAFAYYIEPQHRDVLQLYAPEAFNYMDKFNEQEINLSTRDRIDVLKGEKALKASGLVPEHADAFKKI ARELNTYILFRPVNKLATNLIKSGVATKGLNVHGKSSDWGPVAGYIPFDQDLSKKHGQQLAVEKGNLENK KSITEHEGEIGKIPLKLDHLRIEELKENGIILKGKKEIDNGKKYYLLESNNQVYEFRISDENNEVQYKTKEGKI TVLGEKFNWRNIEVMAKNVEGVLKPLTADYDLFALAPSLTEIKKQIPQKEWDKVVNTPNSLEKQKGVTNL LIKYGIERKPDSTKGTLSNWQKQMLDRLNEAVKYTGYTGGGCGNHGTEQDNEEFPEKDNEIFIINPEGEFIL TKNWEMTGRFIEKNITGKDYLYYFNRSYNKIAPGNKAYIEWTDPITKAKINTIPTSAEFIKNLSSIRRSSNVG VYKDSGDKDEFAKKESVKKIAGYLSDYYNSANHIFSQEKKRKISIFRGIQAYNEIENVLKSKQIAPEYKNYF QYLKERITNQVQLLLTHQKSNIEFKLLYKQLNFTENETDNFEVFQKIIDEK

**LF_N_-LC/A^C699S^** AGGHGDVGMHVKEKEKNKDENKRKDEERNKTQEEHLKEIMKHIVKIEVKGEEAVKKEAAEKLLEKVPSD VLEMYKAIGGKIYIVDGDITKHISLEALSEDKKKIKDIYGKDALLHEHYVYAKEGYEPVLVIQSSEDYVENT EKALNVYYEIGKILSRDILSKINQPYQKFLDVLNTIKNASDSDGQDLLFTNQLKEHPTDFSVEFLEQNSNEVQ EVFAKAFAYYIEPQHRDVLQLYAPEAFNYMDKFNEQEINLSLEELKDQGGSGGSMPFVNKQFNYKDPVNG VDIAYIKIPNAGQMQPVKAFKIHNKIWVIPERDTFTNPEEGDLNPPPEAKQVPVSYYDSTYLSTDNEKDNYL KGVTKLFERIYSTDLGRMLLTSIVRGIPFWGGSTIDTELKVIDTNCINVIQPDGSYRSEELNLVIIGPSADIIQFE CKSFGHEVLNLTRNGYGSTQYIRFSPDFTFGFEESLEVDTNPLLGAGKFATDPAVTLAHELIHAGHRLYGIAI NPNRVFKVNTNAYYEMSGLEVSFEELRTFGGHDAKFIDSLQENEFRLYYYNKFKDIASTLNKAKSIVGTTA SLQYMKNVFKEKYLLSEDTSGKFSVDKLKFDKLYKMLTEIYTEDNFVKFFKVLNRKTYLNFDKAVFKINIV PKVNYTIYDGFNLRNTNLAANFNGQNTEINNMNFTKLKNFTGLFEFYKLLCVRGIITSKTKSLDKGYNKEN LYFQ

## ACKNOWLEDGEMENTS

The authors thank Kaitlin Goldstein, Sebastien Sannajust, Michelle Heyang and Nicholas LoRocco for excellent technical assistance and Tiphaine Voisin for helpful discussions.

## AUTHOR CONTRIBUTIONS

N.J.Y., R.J.C. and I.M.C. conceived the project. N.J.Y., D.N. and A.K-C. performed experiments and data analysis. J.I., A.B. and T.H. performed HCS microscopy analysis and quantified DRG and spinal cord sections. H.X.B.Z. and B.P.B. performed electrophysiological analysis. P.R., A.N. and O.B. provided human iPSC derived nociceptors. J.M.P. provided *B. anthracis* strains. M.L., B.L.P. and S.H.L. provided native and chimeric anthrax toxins. S.M.L., S.P., V.T. and K.A.F. purified and characterized chimeric anthrax-botulinum toxin. N.J.Y., J.I. and I.M.C. wrote the manuscript with input from all authors.

## DATA AND MATERIALS AVAILABILITY

All data associated with this study are present in the main text or the Supplementary Materials. The LFN-LC/A^C699S^ construct is available from Ipsen and require a material transfer agreement.

